# Regulation of murine follicle-stimulating hormone β subunit transcription by newly identified enhancers

**DOI:** 10.1101/2025.11.21.689832

**Authors:** Yangfan Jin, Hailey Schultz, Luisina Ongaro, Gauthier Schang, Xiang Zhou, Carlos Agustin Isidro Alonso, Michel Zamojski, German Nudelman, Natalia Mendelev, Shinsuke Onuma, Corrine K. Welt, Louise M. Bilezikjian, Stuart C. Sealfon, Frederique Ruf-Zamojski, Daniel J. Bernard

## Abstract

Activin-class ligands of the transforming growth factor β family induce follicle-stimulating hormone (FSH) production by pituitary gonadotrope cells in mice via the actions of the transcription factors SMAD3, SMAD4, and FOXL2, which bind to *cis*-elements in the FSHβ subunit (*Fshb*) promoter. An enhancer region for murine *Fshb* transcription was identified *in vitro*. However, deletion of the region using CRISPR-Cas9 did not affect FSH synthesis or secretion in mice. Using single-nucleus ATAC-seq of whole murine pituitaries, we identified three additional open chromatin regions upstream of *Fshb* exclusively in gonadotropes. These regions, as well as the *Fshb* gene, were fully or partially closed in gonadotropes of FSH-deficient mice with genetically or pharmacologically inactivated activin type II receptors. The initially characterized enhancer region did not significantly alter basal or activin-stimulated murine *Fshb* promoter-reporter activity in homologous LβT2 cells. In contrast, the other three open chromatin regions enhanced basal and activin A-stimulated *Fshb* promoter-reporter activity in LβT2 cells, with the two most distal showing the greatest effects. These two regions were open, exhibited enrichment of the enhancer mark H3K27ac, and were bound by SMAD2/3 and FOXL2 in response to activin A in LβT2 cells. The most distal enhancer exhibited strong FOXL2 and weak SMAD4 binding in gel shift assays. SMAD4, but not FOXL2, directly bound the other distal enhancer. Mutation of defined FOXL2 and SMAD4 *cis*-elements diminished enhancer activity in reporter assays in LβT2 cells. Collectively, the data indicate that there may be as many as four activin-sensitive enhancers upstream of murine *Fshb*.

## Introduction

Follicle-stimulating hormone (FSH) is a key regulator of mammalian reproduction, especially in females (1–3). FSH targets ovarian granulosa cells to stimulate follicle growth and estrogen biosynthesis (4,5). In males, FSH promotes Sertoli cell proliferation during development and spermatogenesis in adulthood (6). FSH is a dimeric glycoprotein produced by pituitary gonadotrope cells and is composed of the chorionic gonadotropin α (CGA) and FSHβ subunits (product of the *Fshb* gene) (7). FSHβ synthesis is rate-limiting in the production of the mature hormone (8) and is driven by hypothalamic gonadotropin-releasing hormone (GnRH) and activin-class ligands, including myostatin, in the transforming growth factor β (TGFβ) family (hereafter, activins) (3,9). How GnRH stimulates *Fshb* expression has not been resolved (3,10). In contrast, the mechanisms through which activins regulate *Fshb* promoter activity have been well-delineated *in vitro* (11–13), with some validation *in vivo* in mice (14–16).

Activins bind to activin type II receptors (ACVR2A and ACVR2B), which are transmembrane serine/threonine kinases (17,18). Mice lacking these receptors in their gonadotropes are FSH-deficient, leading to hypogonadism in males and infertility in females due to a block in ovarian follicle growth at the early antral stage (16). Upon ligand binding to the activin type II receptors, type I receptors, such as TGFBR1 and ACVR1B, are recruited into the complex (3,19). Type I receptors are also serine/threonine kinases, which are activated via phosphorylation by type II receptors. Type I receptors then phosphorylate the intracellular signaling proteins, SMADs 2 and 3 (20). SMAD3 is particularly important for FSH synthesis in murine gonadotropes (21,22). Phosphorylated SMAD3 forms a complex with the co-SMAD, SMAD4, and the forkhead box L2 (FOXL2) transcription factor, and binds to the murine and porcine *Fshb* promoters to stimulate transcription (12,23–25). Gonadotrope-specific SMAD3, SMAD4, and FOXL2 knockout mice are FSH deficient, demonstrating the necessity of these proteins for *Fshb* expression *in vivo* (14,15,21,25,26). Loss of SMAD4 and FOXL2 also reduces expression of human *FSHB* in a humanized transgenic mouse model (27). These data suggest that mechanisms of activin-stimulated FSH production may be conserved across species. That said, SMADs and FOXL2 appear to act via both common and species-specific regulatory elements in the *Fshb*/*FSHB* promoter regions investigated to date (12,28,29).

Though activin regulation of *Fshb*/*FSHB* promoters has been studied extensively, previously undiscovered enhancers may also play important roles in FSH synthesis. One open chromatin region 5’ of murine *Fshb* (chr2 107,076,909-107,077,358) is well-conserved between mice and humans, and reportedly exhibits modest enhancer activity in reporter assays in murine gonadotrope-like LβT2 cells (30). The corresponding region in humans (chr11 30,204,583-30,205,051) similarly enhanced *FSHB* promoter-reporter activity in these cells and increased the response to GnRH or activin A (31). Interestingly, two single-nucleotide polymorphisms (rs11031005, rs11031006) associated with reduced FSH levels in women are located within this enhancer region (32,33), though rs11031006 modestly increased *FSHB* promoter activity in reporter assays (30). Collectively, these findings suggest that the conserved open chromatin region acts as a hormone-responsive enhancer of *Fshb*/*FSHB*.

Here, we examined the role of this putative enhancer *in vivo* in knockout mice. In addition, using single-nucleus ATAC-sequencing, we identified three additional regions of co-accessibility upstream of the *Fshb* gene in murine gonadotropes. We then assessed whether these regions possessed enhancer activity *in vitro* and how they may mediate *Fshb* transcriptional regulation by activins.

## Methods

### Mice

To generate mice with the enhancer (Enh 2) knocked out **(Supplementary Fig. S1A** in (34)), two sgRNAs were designed to target chr2: 107,076,832-107,077,562 (GRCm38/mm10), a region of open chromatin ∼17 kb upstream of the *Fshb* gene(30). The DNA repair template was designed to remove the open chromatin region but otherwise leave the locus intact. Mice were generated by electroporation of the two sgRNAs, the repair template (**Supplementary Table S1** in (34)), and Cas9 protein into one-cell C57BL6/N mouse zygotes in the McGill Integrated Core for Animal Modeling (MICAM). The electroporated zygotes were cultured overnight at 37°C in a 5% CO2 incubator. Embryos were then transferred to the oviducts of pseudopregnant females (CD-1 strain). Genomic DNA was extracted from ear biopsies of live pups and then genotyped by PCR using the primers indicated in **Supplementary Table S1** (**Supplementary Fig. S1B**) in (34). Two founder males had the designed 426-base pair (bp) deletion in either one or both alleles. The deletion was germline transmissible (the modified allele is now referred to as Rr466^em1Djb^; MGI 7574263). The two founders were then backcrossed to wild-type C57BL6/N mice for two generations to minimize potential off-target effects.

Mice were housed on a 12-hour light/12-hour dark cycle (lights on at 7 am). Food and water were given *ad libitum*. All animal work was conducted with the approval of the DOW-A Animal Care Committee at McGill University (protocol no. 5204), and in accordance with federal and institutional guidelines.

### Bulk ATAC-seq of LβT2 cells

LβT2b cells (35) were seeded in 12-well plates at 350,000 cells/well in Dulbecco’s Modified Eagle Medium (DMEM, Corning, cat# 10-017-CV)/10% fetal bovine serum (FBS Performance, Wisent Bioproducts, cat#098-150) with six replicates each for quality control (QC) by quantitative real-time polymerase chain reaction (qPCR) and two replicates for bulk ATAC-seq. The next day, cells were switched to serum-free DMEM overnight. Cells were treated the next day either with vehicle or with 1 nM activin A (R&D Systems cat#338-AC) in serum-free DMEM for 6 hours. For qPCR QC, the cells were lysed in RNA lysis buffer before proceeding with RNA extraction, reverse-transcription, and SYBR green qPCR to assess *Fshb* and Ribosomal protein S11 (*Rps11*) as previously described (35,36). Primer sequences were previously published (37). For bulk ATAC-seq, cells were trypsinized, counted, and 50,000 cells were placed per tube before proceeding with the ATAC protocol (38). Dual-side purification of the libraries was performed, and the libraries were resuspended in 20 μl of H2O. All libraries were QCed using Bioanalyzer (Agilent, Santa Clara, CA) and Qubit (fluorometric quantitation; Thermo Fisher Scientific, Waltham, MA). Libraries were sequenced on a Novaseq sequencer at the New York Genome Center. The nf-core/atacseq (v2.1.2) pipeline was utilized to analyze bulk ATAC-seq data (39). Preprocessed reads were aligned to GRCm38 reference genome using bowtie2 (40); the rest of the pipeline was run using default settings. Coverage plots were generated using Signac (v1.14.0) with a window size of 500 (41). These datasets are available at the GEO repository: GSE310458.

### Single nucleus ATAC-seq of murine pituitaries

Pituitary glands were dissected from 8- to 10-week-old wild-type (WT) and Enh 2 knockout (KO), bimagrumab and IgG injected, and control and *Acvr2a/b* double knockout (dKO) males (16). Pituitary glands were snap-frozen and stored at -80°C. Nuclei were isolated on ice from snap-frozen pituitaries as described in (42,43). Briefly, RNAse inhibitor (NEB MO314L) was added to the homogenization buffer [0.32 M sucrose, 0.1 mM EDTA, 10 mM Tris-HCl, pH 7.4, 5 mM CaCl2, 3 mM Mg(Ac)2, 0.1% IGEPAL CA-630], 50% OptiPrep (Sigma; cat# D1556), 35% OptiPrep, and 30% OptiPrep right before isolation. Each pituitary was homogenized in a Dounce glass homogenizer (1 ml, VWR cat# 71000-514), and the homogenate was filtered through a 40 μm cell strainer. An equal volume of 50% OptiPrep was added, and the gradient was centrifuged (SW32 rotor at 10,200 × g; 4°C; 25 min). Nuclei were collected from the interphase, washed, resuspended in 1X nuclei dilution buffer for snATACseq (10x Genomics, Pleasanton, CA) and counted (Revvity Cellometer K2). snATACseq was performed following the Chromium Next GEM Single Cell ATAC Reagent Kits v1.1 User Guide (10x Genomics, Pleasanton, CA). Transposition was performed in 10 μl at 37°C for 60 min on 4,000 to 10,000 nuclei, depending on the samples, before loading the Chromium Chip H (PN-2000180) for GEM generation. Barcoding was performed in the emulsion (12 cycles) following the Chromium protocol. Libraries were indexed for multiplexing (Chromium i7 Sample Index N, Set A kit PN-1000212). Libraries were sequenced on a Novaseq-X sequencer at the Cedars-Sinai Applied Genomics, Computation & Translational Core. snATACseq data were processed using Cell Ranger-ATAC pipeline version 2.2.0.

snATAC-seq data were clustered using Signac (v1.14.0) following standard procedures(41). Cell types were annotated based on the chromatin accessibility of pituitary cell type markers as previously described (42). snATAC-seq datasets were integrated with Seurat (v 5.3.0) (44) with the Harmony (v1.2.3) package (45) following standard procedures for snATAC-seq data. The processed and analyzed murine snATAC-seq dataset collection for **Supplementary Figs. S2A-B** in (34) were previously published (42). Coverage plots were generated using Signac with a window size of 500. These datasets are available at the GEO repository: GSE310460.

### Gonadectomy

Male and female mice were gonadectomized at 8 weeks of age following standard operating procedures 206 and 207 of McGill University. In sham-operated (control) female mice, all the procedures were the same, except that the ovaries were not cauterized. After two weeks recovery from surgery, mice were anesthetized with isoflurane and euthanized by CO2 asphyxiation. Their pituitary glands were collected, snap-frozen in liquid N2, and stored at -80°C. Sham-operated females were randomly cycling at the time of collection in the afternoon. Blood was collected as described below.

### Bimagrumab injections

Murine bimagrumab (RRID:AB_3718172) was expressed by transient transfection in ExpiCHO-S™ cells, using the ExpiCHO™ Expression System kit (Gibco #A29133) following the manufacturer’s guidelines. Clarified harvest was then purified by Protein A affinity chromatography and the purified antibody was formulated in D-PBS. The final formulation was stored at -80°C until use. Nine-week-old male C57BL6/N mice were injected with a single dose of 20 mg/kg of mouse IgG (Sigma Aldrich I5381; RRID:AB_1163670) or bimagrumab subcutaneously. Submandibular blood was collected before injections. The animals were euthanized 3 days after antibody injections, and blood was collected by cardiac puncture to measure FSH levels. Pituitary glands were flash-frozen in liquid N2 and subjected to snATAC-seq.

### Blood collection and hormone analyses

In most cases, blood was collected by cardiac puncture in the afternoon, at the time of sacrifice, from 8- to 10-week-old males and females. Females that were not subjected to surgery were euthanized at 8-9 weeks of age at 7:00 am on the morning of estrus (determined by vaginal cytology). Submandibular blood was collected from 8-week-old males prior to castration and then by cardiac puncture 2 weeks after castration. Blood was allowed to coagulate at room temperature for 30 min and was then centrifuged for 10 min at 3000 rpm. Serum was collected and stored at -20°C. Whole blood was also collected from tails of pre- and post-gonadectomized mice, diluted 1/30 in PBS with 5% Tween-20 (PBS-T), snap-frozen in liquid N2, and stored at - 80°C.

Serum FSH, serum LH, and whole blood LH levels were determined using validated in-house enzyme-linked immunosorbent assays (ELISAs), as described previously (46,47). For the FSH ELISA, guinea pig anti-mouse FSH capture antibody (AFP-1760191, RRID: AB_2665512) and rabbit anti-rat FSH antibody RIA (NIDDK-anti-rFSH-S-11, AFP-C0972881, RRID: AB_2687903) for detection were used. For the LH ELISA, we used a mouse monoclonal anti-bovine LHβ subunit antibody (RRID: AB_2756886) for capture and a rabbit polyclonal anti-rat LH antiserum (AFP240580Rb, RRID: AB_2665533) for detection.

### Tissue collection

Pituitaries were collected from WT and KO mice, snap-frozen in liquid N2, and stored at - 80°C. Testes and seminal vesicles from WT and KO mice were dissected and weighed. For mice that were not subjected to surgery, WT and KO females were euthanized at 9-10 weeks of age at 7:00 am on the morning of estrus (determined by vaginal cytology). Their ovaries and uteri were dissected and weighed.

### Reverse transcription and quantitative PCR

RNA was extracted from homogenized pituitaries using TRIzol reagent (15596018, Invitrogen, Waltham, MA, USA) following the manufacturer’s protocol. Concentration of RNA was determined by NanoDrop spectrophotometry. For all experiments, 200 ng of total RNA were treated with RQ1 DNase and then reverse-transcribed into cDNA using random hexamer primers (C1181, Promega) and Moloney murine leukemia virus reverse transcriptase (M1701, Promega). Each reverse transcription reaction was diluted 1:2 in DEPC-treated H2O before qPCR analyses. PCR was performed using EvaGreen (ABMMmix, Diamed, Missisauga, ON, Canada) with primers listed in **Supplementary Table S1** in (34) using a Corbett Rotorgene 600 instrument (Corbett Life Science, Sydney, NSW, Australia). All primers were validated for efficiency and specificity. Relative mRNA levels were determined using the 2^-ΔΔCt^ method. Gene expression was normalized to ribosomal protein L19 (*Rpl19*).

### DNA constructs

The wild-type -1990/+1 murine *Fshb*-luc promoter-reporter in pGL3-Basic was described previously (20). Regions of open chromatin upstream of *Fshb*, hereafter Enh 1-4 (Enh 1: Chr2 107,072,061-107,073,651; Enh 2: Chr2 107,076,912-107,077,479; Enh 3: Chr2 107,118,588-107,119,297; Enh 4: Chr2 107,126,502-107,127,575 GRCm38/mm10) were amplified by nested PCR from C57BL6/N mouse genomic DNA extracted from tails. Inner primers for each enhancer region were used to add *Sal*I restriction sites at both ends (**Supplementary Table S1** in (34)). Each enhancer region was ligated into the *Sal*I site in the murine -1990/+1 *Fshb*-luc reporter plasmid, downstream of the luciferase reporter gene. Plasmids with insertions in both orientations were obtained. Enh 1, 3, and 4 were also ligated in both orientations into the same site in the pGL3-Promoter vector (Promega), which contains the SV40 promoter. Truncations of Enh 3 and 4 were produced by PCR and similarly ligated (**Supplementary Table S1** in (34)).

The FLAG-FOXL2 expression vector was described previously (29).

GST-SMAD3-MH1 (codons 1-200), GST-SMAD4-MH1 (codons 1-229) in pGEX2TK vector were described in (48). GST-FOXL2-FHD (forkhead domain, codons 48-144) was prepared by PCR amplifying the needed sequence from the FLAG-FOXL2 construct with the primers described in (**Supplementary Table S1** in (34)) and ligating the fragment in-frame into the *BamH*I/*Xho*I sites in the pGEX4T vector.

All mutant promoter/enhancer reporters were generated using the QuikChange protocol and primers in **Supplementary Table S1** in (34). All constructs were verified using Sanger sequencing (Genome Quebec).

### Cell lines

Human embryonic kidney (HEK) 293T cells (ATCC CRL-3216; RRID:CVCL 0063; provided by Dr. Terry Hébert, McGill University) were cultured in Dulbecco’s modified Eagle’s medium (DMEM, 319-005-CL, Wisent, St-Bruno, QC, Canada) containing 5% (v/v) fetal bovine serum (098150, Wisent). Immortalized murine gonadotrope-like LβT2 (LβT2b cells (35) used for bulk ATAC-seq and ChIP-seq) and αT3-1 cells (RRID:CVCL 0398 for LβT2, CVCL0149 for αT3-1) provided by Dr. Pamela Mellon (University of California, San Diego, CA, USA) were cultured in DMEM with 10% (v/v) fetal bovine serum. All cells were cultured at 37°C with 5% CO2 in a humidified incubator.

We authenticated the cell lines used in this study as described in (49). Frozen aliquots of each LβT2 line were shipped to Idexx BioResearch (Columbia, MO) for cell line authentication. The *CellCheck Mouse Plus* profile performed by Idexx included (i) cell line identification by STR DNA profiling (**Supplementary Table S2** in (34)), mycoplasma testing, and (ii) multiplex PCR-based interspecies contamination check for the mouse, rat, human, Chinese hamster, and African green monkey. STR profiling was performed for 19 species-specific STR markers. All three gonadotrope cell lines in this study were negative for mycoplasma and confirmed of mouse origin.

The LβT2 line used for reporter assays and bulk ATAC (LβT2b) in this manuscript differed from the two LβT2 lines used for ChIP-seq assays (LβT2-FOXL2 and LβT2-GFP) only by marker MCA-7-1, highlighted in **Supplementary Table S2** in (34).

### Promoter-reporter assays

Promoter-reporter assays were performed as previously described (50). Briefly, LβT2b cells were seeded at a density of 150,000 cells per well in 48-well plates. The next day, the cells were transfected with 225 ng/well of the indicated reporter plasmid constructs using Lipofectamine 3000 (L3000015, ThermoFisher Scientific, Burlington, ON, Canada) following the manufacturer’s protocol. Twenty-four hours after transfection, cells were serum starved overnight. The next day, cells were either treated with 1, 0.5, 0.25, 0.125, 0.0625, 0.05, 0.03125 or 0.0015625 nM activin A (388-AC-050, R&D Systems, Minneapolis, MN, USA) for 6 h, 10 µM SB431542 (S4317, Sigma-Aldrich) for 24 h, or serum-free media as control for activin A, and serum-free media with DMSO as control for SB431542. Cells were lysed using 50 µL/well passive lysis buffer (25 mM Tris-phosphate [pH 7.8], 10% [v/v] glycerol, 1% [v/v] Triton X-100, 1 mg/mL bovine serum albumin, 2 mM ethylenediaminetetraacetic acid [EDTA]) for 10 minutes at room temperature with agitation. One-hundred µL of assay buffer (15 mM potassium phosphate [pH 7.8], 25 mM glycylglycine, 15 mM MgSO_4_, 4 mM EDTA, 2 mM adenosine triphosphate, 1 mM dithiothreitol, 0.04 mM D-luciferin) were added to 20 µL of cell lysis supernatant, and luciferase activity measured using an Orion II microplate luminometer (Berthold Detection Systems, Oak Ridge, TN, USA). Experiments were performed in either technical duplicates or triplicates, and the experiments were repeated at least three times, as indicated in the figures or figure legends.

### Chromatin immunoprecipitation sequencing

LβT2 cells stably expressing FLAG-tagged FOXL2 (FLAG-mFOXL2) were generated by lentivirus transduction of a cassette (pSpike14) that permits constitutive GFP expression and tetracycline/doxycycline (DOX)-inducible, IRES-mediated tandem expression of a fluorescent reporter (Neptune) and a protein of interest (i.e., FLAG-mFOXL2 or vector only), as published in (51) (the pSpike14 vector was designed and provided by Drs. Geoffrey Wahl and Benjamin Spike, Salk Institute). Single cell suspensions of transduced LβT2 (2×10^7^ cells) were FACS sorted, and GFP-positive cells that were recovered (0.5×10^6^ cells) were seeded on 10 cm tissue culture dishes for expansion and storage of aliquots of pooled GFP-positive cells in liquid N2 for further evaluation. The cells were subsequently functionally validated to express DOX-inducible FLAG-mFOXL2 by western blot analysis, and retain characteristics of parental LβT2 cells, including activin-responsiveness in luciferase experiments (data not shown).

For ChIP-seq experiments, stably transduced LβT2-FLAG-mFOXL2 cells were seeded in 15 cm tissue culture plates in DMEM supplemented with 10% FBS (Tet system approved FBS, Clontech No 631106). After a recovery period of 24 h, DOX-induction was initiated in fresh medium by the addition of DOX to a final concentration of 2% (w/v) for 48 h. After 48 h, the cells were washed and equilibrated for 2 h in DMEM supplemented with 2% FBS before adding vehicle or 1 nM activin A for 1 h. Cross-linking of chromatin-protein complexes was achieved by the addition of a formaldehyde solution directly to the culture medium to a final concentration of 1%. The experimental strategy to prepare cross-linked complexes suitable for ChIP studies were the same as previously published (52). Duplicate samples for each of the vehicle or activin A treatment group consisted of pooled samples from three 15 cm plates. The enrichment of SMAD2/3-, FLAG-FOXL2- and H3K27ac-bound complexes were achieved essentially as previously described (52). SMAD complexes were enriched using an IgG-purified rabbit anti-SMAD2/3 antibody previously characterized for its specificity and use for ChIP experiments (generated in the laboratory of Dr. Wylie Vale at the Salk Institute; RRID:AB_3718178) (52).

The M2-anti-FLAG MAb (Sigma-Aldrich, F1804, RRID:AB_262044) was used to enrich FLAG-mFOXL2-bound complexes and anti-H3K27ac (Abcam, ab4729, RRID: AB_2118291) was used to mark open chromatin regions. Antibody-bound complexes were enriched with Protein-A (Abcam, ab214288) or Protein-G magnetic DYNA beads (Abcam, ab286842).

Sequencing libraries appropriate for Illumina platforms were prepared using NEXTFlex DNA sequencing kits (Bioo Scientific, Austin, TX). Briefly, ChIP DNA fragments were blunted using the Klenow fragment followed by the addition of 3’-overhangs and ligation of barcoded adapters, with intervening cleanup steps with magnetic beads/PEG. After PCR amplification with adapter-specific primers, 300-400 bp fragments were size-selected by differential PEG precipitation onto magnetic beads. ChIP-seq libraries were sequenced for 51 cycles to a depth of 20 million reads using single-end 50 bp reads on an Illumina HiSeq2500 sequencer (at the Ravazi Newman Integrative Genomic and Bioinformatics Core of the Salk Institute for Biological Studies). The nf-core/chipseq pipeline (v2.0.0) was utilized to analyze ChIP-seq data

(39). Preprocessed reads were aligned to the GRCm38 reference genome using bowtie2 (40); the rest of the pipeline was run using default settings. Coverage plots were generated using Signac (v1.14.0) with a window size of 500(41). De novo motif analysis for enriched regulatory elements and treatment-related changes were additionally performed using the software suite, HOMER by the Bioinformatics Core of the Salk Institute for Biological Studies (53). These datasets are available at the GEO repository: GSE310459.

### Electrophoretic mobility shift assays

HEK293T cells were seeded at a density of 3×10^6^ in 10-cm plates for electrophoretic mobility shift assays (EMSA). The next day, cells were transfected with 7 µg FLAG-FOXL2 or empty vector (pcDNA3.0) using polyethylenimine (PEI) at a ratio of 1:3 for 2 h and then changed to media with 5% FBS. Cells were starved in serum-free DMEM for 24 hours the next day. Cells were washed twice with PBS, collected in 1 mL PBS using a cell scraper, and centrifuged for 5 min at 400 x g. Supernatant was removed, 500 µL 1x lysis buffer (10 mM HEPES pH 7.9, 1.5 mM MgCl2, 10 mM KCl, 10 µM leupeptin, pepstatin A, and aprotinin, 1 mM PMSF) were added to the pellet, cells were resuspended, and incubated for 15 min on ice.

Sixteen µL of NP-40 was added to the swollen cells in lysis buffer to reach a final concentration of 0.3%, and the tube was vortexed vigorously for 10 sec. Samples were immediately centrifuged for 30 sec at 11,000 x g, supernatant was removed, and crude nuclei pellet was resuspended in 100 µL of complete cell extraction buffer (20 mM HEPES pH 7.9, 1.5 mM MgCl2, 0.42 M NaCl, 0.2 mM EDTA, 25% glycerol, 10 µM leupeptin, pepstatin A, and aprotinin, 1 mM PMSF) and was agitated using a vortex at medium to high speed for 15-30 min at 4°C. Nuclear extracts were obtained from the supernatant after centrifugation at 21,000 x g for 5 min at 4°C, snap-frozen in liquid N2, and stored at -80°C. Protein concentrations were determined by Bradford assay (BioRad).

Recombinant GST-SMAD3-MH1, GST-SMAD4-MH1, and GST-FOXL2-FHD were prepared following protocols adapted from (11,54,55). In brief, BL21 cells transformed with GST-SMAD3-MH1, GST-SMAD4-MH1, or GST-FOXL2-FHD plasmids were grown in 1L LB containing 100 µg/ml ampicillin while shaking at 250 rpm at 37°C until OD600 was 0.5-0.7. Protein expression was induced by adding IPTG (Sigma-Aldrich, AM9462) to a final concentration of 0.1 mM and incubating at 37°C while shaking for 4 h. Cells were harvested and resuspended in lysis buffer (1X PBS, 0.1 mM PMSF, and 0.1 mM protease inhibitor cocktail IV, Calbiochem), then lysed by sonication (Misonix Sonicator 3000, 15-sec on, 45-sec off, 45% input, 6 min total time on) at 4°C. Triton X-100 was added to the mixture to a final concentration of 1%. After mixing for 30 min at 4°C, the mixtures were centrifuged at 10,000xg for 10 min at 4°C, and supernatant combined with 0.75 mL pre-washed Glutathione Sepharose beads (Sigma-Aldrich GE17-0756-01). Together, beads and protein mixture were incubated for 1 h at 4°C with rotation. Samples were then centrifuged at 500xg for 3 min at 4°C, and beads washed four times with wash buffer (1X PBS + 1% Triton X-100). Beads were transferred into Bio-Rad Chromatography Columns (Econo-Column, #7372512) and washed twice with cold 1X PBS. Proteins were then eluted using 20 mM reduced glutathione, snap-frozen in liquid nitrogen, and stored at -80°C. Before proceeding to EMSA, proteins were run on SDS-PAGE gels to determine their purity. Protein concentrations were determined by Bradford assay (BioRad).

For FOXL2 gel shifts, 10 µg of nuclear extract or 800 ng recombinant protein were incubated with or without unlabeled competitor probes (**Supplementary Table S1** in (34)), 10x, 100x, 250x molar excess) in 150 mM KCl, 25 mM HEPES (pH 7.2), 5 mM dithiothreitol, 12.5% glycerol, and 500 ng salmon sperm DNA in a final volume of 20 µL at room temperature for 10 min. One-hundred fmol biotin-labeled Enh 4 probe spanning the composite FOXL2/SMAD binding site (**Supplementary Table S1** in (34)) was then added and incubation proceeded for another 20 min at room temperature. Reactions were then run on 5% polyacrylamide gels (44:0.8 acrylamide:bis-acrylamide) in 40 mM Tris-HCl/195 mM glycine (pH 8.5) at 100 volts for 55 min at room temperature. DNA was transferred from the gels onto nylon membranes (GeneScreen Plus) in precooled 40 mM Tris-HCl/195 mM glycine (pH 8.5) at 100 volts for 1 h. Transferred DNA was crosslinked to the membranes using a Stratalinker 1800 (Stratagene, Marshall Scientific) for 15 min. Biotin-labeled DNA was detected by chemiluminescence (LightShift Chemiluminescent EMSA Kit, Thermo Scientific) on an Amersham Imager 600 (GE Healthcare, Chicago, IL, USA).

For gel shifts with GST-SMAD3-MH1 and GST-SMAD4-MH1, 800 ng of recombinant protein or GST alone were incubated with or without unlabeled competitor probes (**Supplementary Table S1,** 10x, 100x, 250x relative to promoter probe, or 50x, 500x, 2500x relative to Enh 3 probe) in 25 mM Tris-HCl (pH 7.5), 80 mM NaCl, 35 mM KCl, 5 mM MgCl2, 1 mM DTT, 10% glycerol and 500 ng salmon sperm DNA in a final volume of 20 µL for 10 min. One-hundred fmol biotin-labeled control *Fshb* promoter probe spanning -132/-92 relative to the transcription start site or 10 fmol biotin-labeled Enh 3 probe (**Supplementary Table S1** in (34)) was then added and incubated for another 20 min at room temperature. Reactions were then run on 5% polyacrylamide gels (29:1 acrylamide:bis-acrylamide) in 0.5X TBE (0.0515 M Tris, 22.5 mM boric acid, 1.25 mM EDTA) for 50 min at room temperature at 100 volts. DNA was transferred from the gels onto nylon membranes (GeneScreen Plus) in precooled 0.5X TBE at 100 volts for 1 h. Transferred DNA was auto-crosslinked to the membranes on a Stratalinker 1800 (Stratagene, Marshall Scientific). Biotin-labeled DNA was detected by chemiluminescence (LightShift Chemiluminescent EMSA Kit, Thermo Scientific) on an Amersham Imager 600 (GE Healthcare, Chicago, IL, USA).

Band intensities were quantified using Image J (NIH, Bethesda, MD, USA) using the Gel Analysis function. For each experiment, signal intensities were normalized to the condition with maximal binding (e.g., protein plus probe without competitors).

### Statistical analyses

Effects of genotype were analyzed using unpaired t-tests or one-way analysis of variance (ANOVA). Luciferase assay data in LβT2 and αT3-1 cells were log-transformed before analysis by two-way ANOVA, followed by Holm-Šidák test. EMSAs were analyzed using one-way ANOVA followed by Tukey’s post hoc tests. Statistical analyses were performed using Prism 8, GraphPad. Alpha was set to P < 0.05.

## Results

### FSH production is not altered in enhancer knockout mice

We used CRISPR-Cas9 with homology-directed repair in murine zygotes to delete a 426-bp region on Chr. 2 (Chr2 107,076,832-107,077,562) containing the putative *Fshb* enhancer (**Supplemental Figs. S1A-B** in (34)). Serum FSH levels did not differ between adult wild-type (WT) and enhancer knockout (KO) mice in either sex (**Figs. 1A-B**). Unexpectedly, serum LH levels were modestly decreased in KO males, but not females (**Figs. 1C-D**). Pituitary *Fshb*, *Lhb*, *Cga*, and *Gnrhr* mRNA levels were not altered in KOs compared to WT males (**Fig. 1E**) or females (**Fig. 1F**). In KO males, testis weights were not altered, but there was a significant decrease in seminal vesicle weights compared to WT (**Fig. 1G-H**). In females, there were no significant differences in ovary or uterus weights between genotypes (**Fig. 1I-J**).

**Figure 1.**
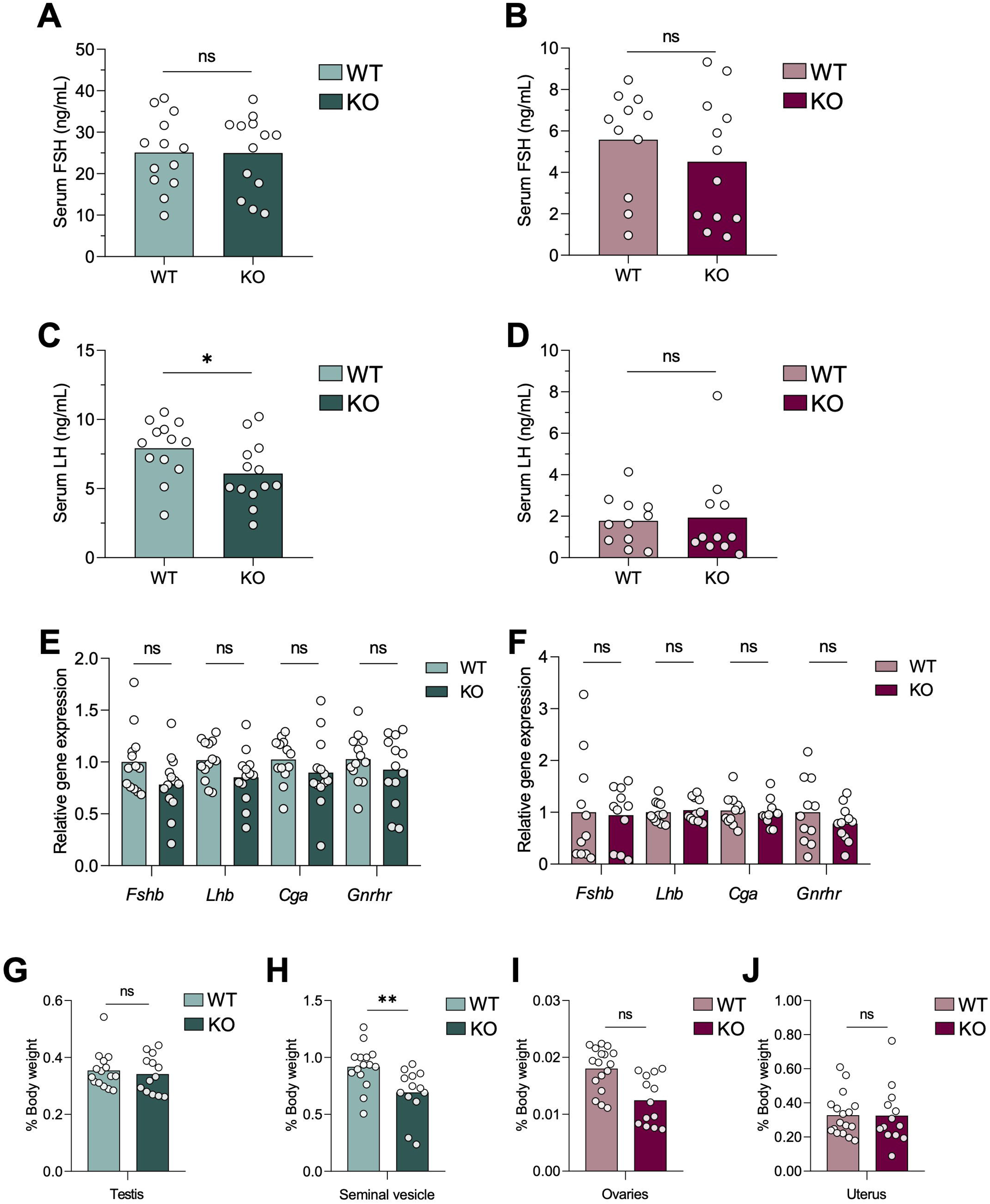
FSH secretion and synthesis are unaltered in enhancer knockout (KO) mice. Serum FSH levels in WT and KO (A) males and (B) females. Serum LH levels in WT and KO (C) males and (D) females. Pituitary *Fshb*, *Lhb*, *Cga,* and *Gnrhr* mRNA expression assessed by RT-qPCR in WT and KO (E) males and (F) females. (G) Testis and (H) seminal vesicle weights of male WT and KO mice. (I) Ovary and (J) uterus weights of female WT and KO mice. Organ weights are normalized to each animal’s body weight. t-tests were performed for statistical analyses, *P < 0.05. **P<0.01. ns, not significant.

To rule out potential masking effects of gonadal hormones (e.g., steroids or inhibins), we also examined gonadectomized WT and KO mice. In males, we employed a within-subjects design and collected blood before and after castration in the same animals. Serum FSH and whole blood LH levels increased post-castration but did not differ between genotypes (**Figs. 2A-B**). Pituitary *Fshb*, *Lhb*, *Cga*, and *Gnrhr* mRNA levels did not differ between genotypes post-castration (**Figs. 2C**).

**Figure 2.**
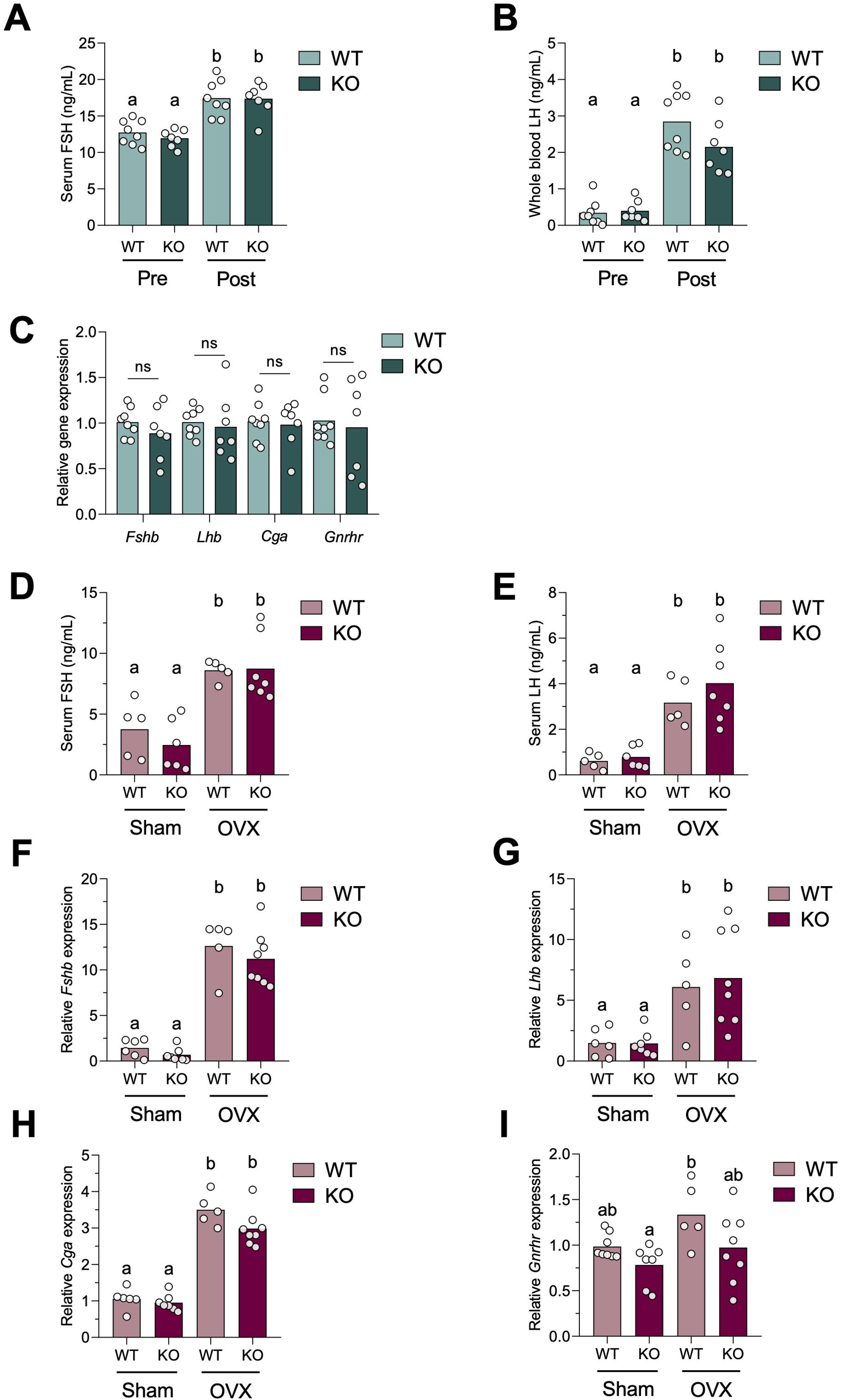
FSH synthesis and secretion are unaltered in castrated or ovariectomized enhancer knockout mice compared to wild type. (A) Serum FSH and (B) whole blood LH in WT and KO males before (pre) and after (post) castration. Two-way ANOVA followed by Holm-Šidák test was performed for statistical analysis. Bars with different letters differ significantly from each other. (C) Pituitary *Fshb*, *Lhb*, *Cga*, and *Gnrhr* mRNA expression assessed by RT-qPCR in male WT and enhancer KO mice after castration. t-tests were performed for statistical analyses. ns, not significant. (D) Serum FSH and (E) whole blood LH in sham (left two bars) and ovariectomized (OVX, right two bars) in WT and KO females. (F) Pituitary *Fshb*, (G) *Lhb*, (H) *Cga,* and (I) *Gnrhr* mRNA expression assessed by RT-qPCR in sham and OVX WT or enhancer KO females. Data were analyzed as in (A).

In females, we employed a between-subjects design comparing ovary-intact (sham) and ovariectomized (OVX) mice of both genotypes. As expected, OVX females had higher serum FSH and whole blood LH levels compared to sham females, but the levels did not differ between WT and KO mice (**Figs. 2D-E**). Pituitary *Fshb*, *Lhb*, *Cga*, and *Gnrhr* expression also did not differ between genotypes, though we observed the expected increases in *Fshb, Lhb,* and *Cga* in OVX animals (**Figs. 2F-I**).

### Identification of three novel *Fshb* enhancers

Enhancer redundancy has been described elsewhere (56,57). To investigate whether there may be other regulatory regions (enhancers) for *Fshb*, we scrutinized our previously published single nucleus (sn) assay for transposase-accessible chromatin using sequencing (snATAC-seq) data from adult WT C57BL6/J male and female mouse pituitaries (42). We identified four open chromatin regions 5’ of *Fshb* that were present in gonadotropes but not in other pituitary cell types (**Fig. 3A(a), Supplemental Figs. S2A-B** in (34)); N.B. 5’ is to the right in these figures).

**Figure 3.**
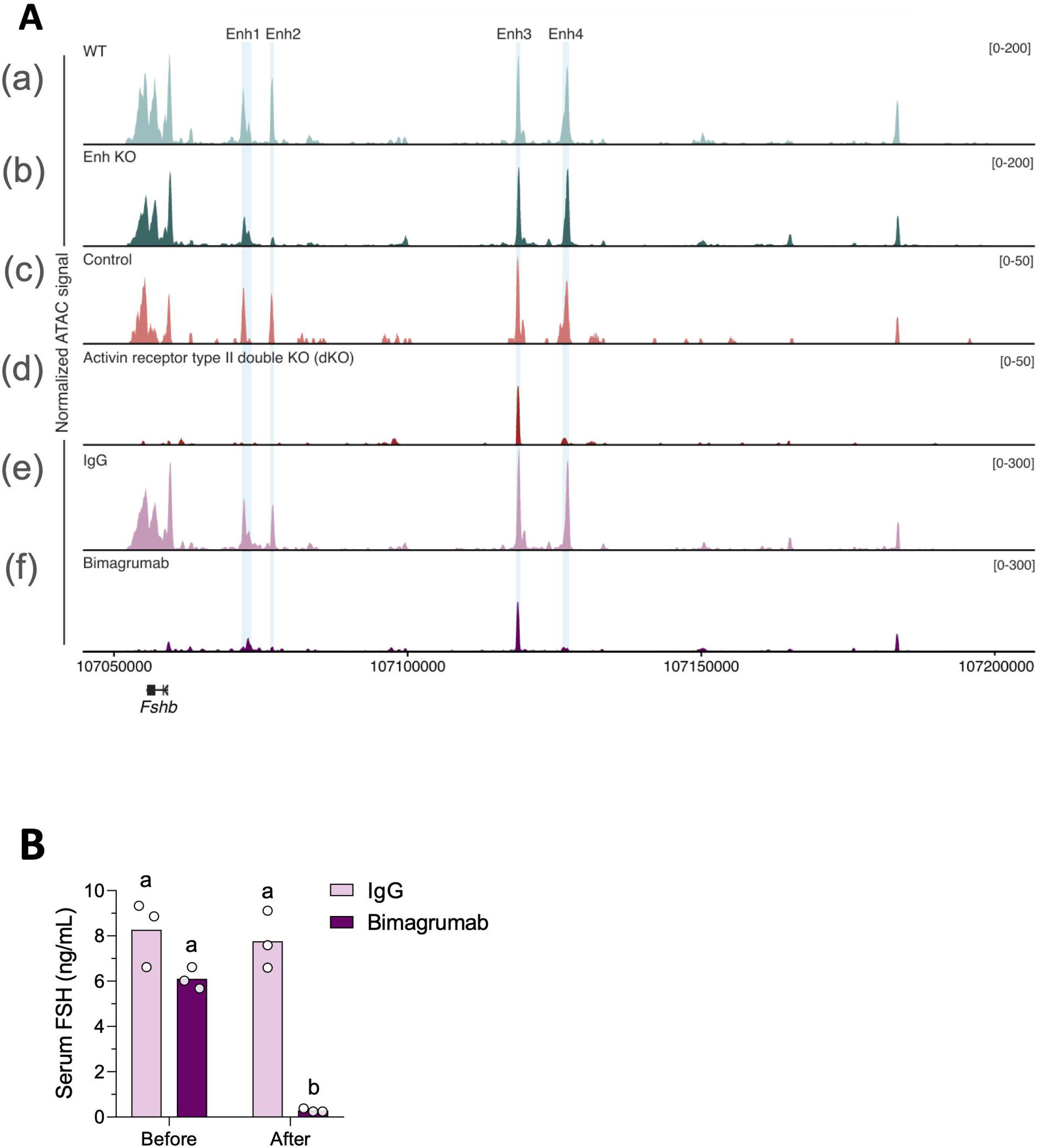
Four upstream enhancers identified by snATAC-seq. (A) Chromatin accessibility of the *Fshb* gene and 5’ flanking sequence (to the right) in gonadotropes of adult male (a) WT or (b) Enh 2 KO mice, (c) control or (d) activin receptor type II double (*Acvr2a/b*) knockout mice, (e) adult mice injected with IgG or (f) Bimagrumab, as revealed using snATAC-seq. (B) Serum FSH levels of adult male mice before and 3 days after injection with IgG (pink bars) or Bimagrumab (magenta bars). Data were analyzed as described in Fig. 2A.

We refer to these regions as enhancers (Enh) 1 to 4, with Enh 1 being the most proximal to the *Fshb* gene, and Enh 4 being the most distal (Enh 1: Chr2 107,072,061-107,073,651; Enh 2: Chr2 107,076,912-107,077,479; Enh 3: Chr2 107,118,588-107,119,297; Enh 4: Chr2 107,126,457-107,127,626, GRCm38/mm10). The enhancer analyzed in the KO mice above corresponds to Enh 2. In snATAC-seq of the pituitaries from KOs mice, there was the expected loss of sequence reads in Enh 2 (**Fig. 3A(b)**). The accessibility of the *Fshb* gene, Enh 3, and Enh 4 was similar compared to WT mice, whereas accessibility of Enh 1 was slightly reduced (**Fig. 3A(b)**).

We previously reported that gonadotrope-specific activin type II receptor (*Acvr2a* and *Acvr2b*) knockout mice do not produce FSH due to the loss of *Fshb* expression (16). We performed snATAC-seq on pituitaries from male *Acvr2a/b* conditional double knockout (dKO) mice and age-matched controls (floxed alleles only). Remarkably, the *Fshb* gene as well as Enh 1, 2, and 4 were completely inaccessible in gonadotropes of dKO mice (**Fig. 3A(c-d)**). In contrast, Enh 3 remained partially open in dKOs. Injection of bimagrumab, an activin type II receptor neutralizing antibody(58), into adult male mice resulted in a 95% reduction in FSH levels (**Fig. 3B**). snATAC-seq on pituitaries from these mice revealed a pattern of chromatin accessibility similar to that of *Acvr2a/b* knockouts, with Enh 1, 2, and 4 completely closed, and Enh 3 remaining somewhat accessible (**Fig. 3A(e-f)**).

### Enh 1, 3, and 4 increase basal and activin A-stimulated *Fshb* promoter activity

To test whether Enh 1-4 possess enhancer activity *in vitro*, we transfected LβT2 cells with a murine -1990/+1 *Fshb*-luciferase promoter-reporter with or without the putative enhancers ligated downstream of the luciferase coding sequence. Enh 1 significantly increased basal and activin A-stimulated *Fshb* promoter activity (**Fig. 4A**). In contrast to an earlier report (30), Enh 2 did not significantly alter basal or activin A-stimulated reporter activity in either orientation (**Fig. 4B**). Enh 3 or Enh 4 increased both basal and activin A-stimulated *Fshb* promoter-reporter activity, with Enh 4 having the quantitatively largest effects (**Figs. 4C-D**). Enh 1, 3, and 4 were active regardless of orientation. When adjusting for changes in basal activity, none of the enhancers significantly increased the fold-induction by 1 nM activin A relative to promoter alone.

**Figure 4.**
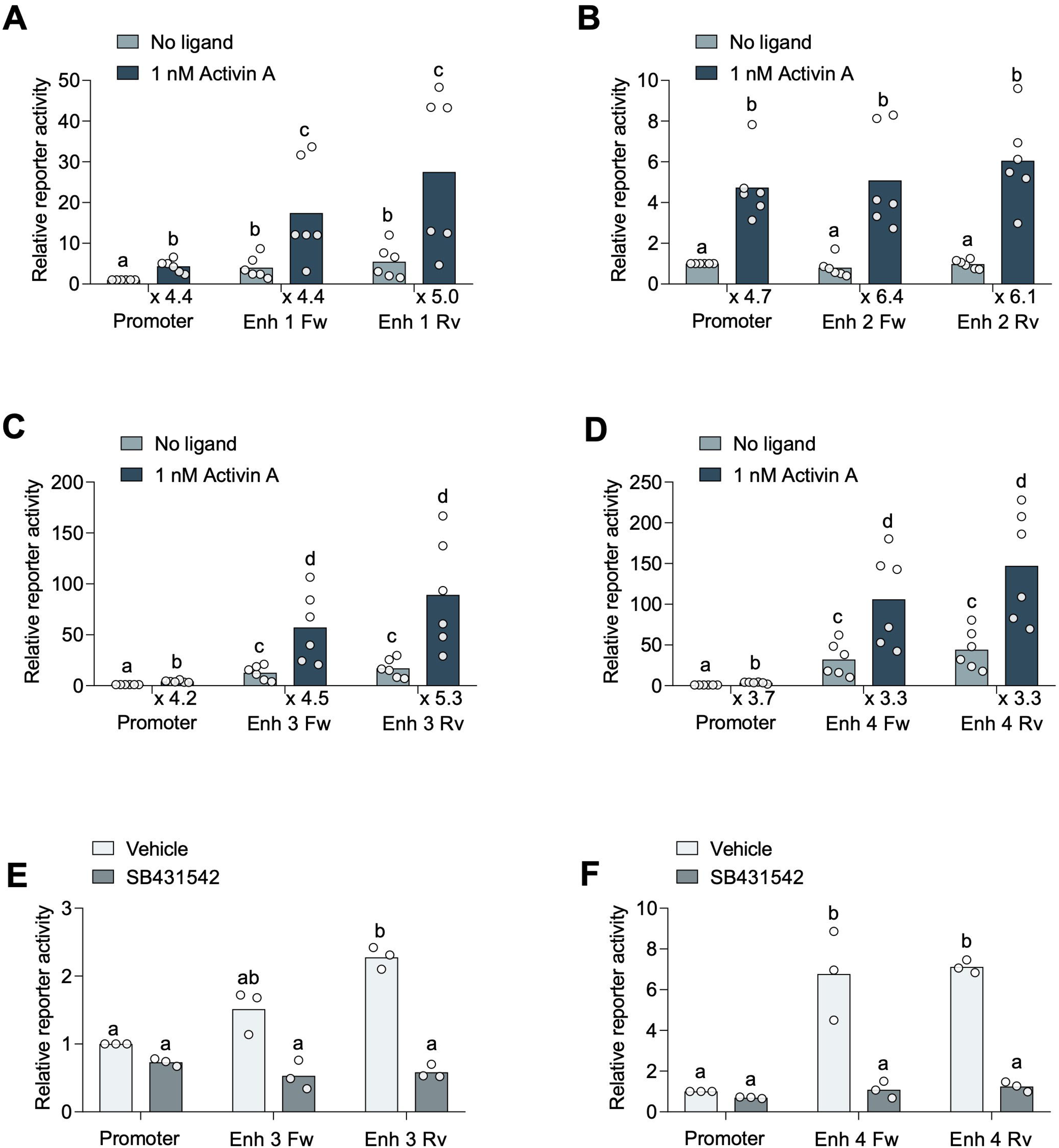
Three of four open chromatin regions exhibit enhancer activity in reporter assays. LβT2 cells were transiently transfected with a -1990/+1 murine *Fshb* promoter-reporter without or with (A) Enh 1, (B) Enh 2, (C) Enh 3, or (D) Enh 4 inserted in either forward (Fw) or reverse (Rv) orientation downstream of luciferase. Cells were treated with no ligand (faint bars) or 1 nM activin A (darker bars) for 6h. Fold-induction by activin A is shown below the x-axis. LβT2 cells were transfected with the -1990/+1 *Fshb* promoter-reporter without or with (E) Enh 3 or (F) Enh 4 inserted in either orientation. Cells were treated with DMSO vehicle (faint bars) or 10 µM SB431542 (dark bars) overnight. Individual points are independent experiments. Data were log-transformed and analyzed by two-way ANOVA followed by Holm-Šidák test. Bars with different letters differ significantly.

Basal *Fshb* mRNA expression in LβT2 cells depends on endogenous activin-like activity (59,60). Here, we similarly observed a reduction in the basal reporter activity conferred by Enh 3 or Enh 4 in cells treated with the activin type I receptor inhibitor, SB431542 (61) (**Figs. 4E-F**).

We did not examine the effects of this inhibitor on Enh 1, given its modest effects on basal activity relative to Enh 3 or Enh 4.

### Enh 1, 3, and 4 increase basal activity of a heterologous promoter

Enh 1, 3, and 4, but not Enh 2, exhibited enhancer activity on the murine *Fshb* promoter in homologous LβT2 cells. We next investigated whether their actions were promoter-specific.

We ligated Enh 1, 3, or 4 into a heterologous promoter-reporter, SV40-luciferase. When transfected into LβT2 cells, all three enhancers increased basal SV40 promoter-reporter activity, but none conferred activin A responsiveness (**Figs. 5A-C**).

**Figure 5.**
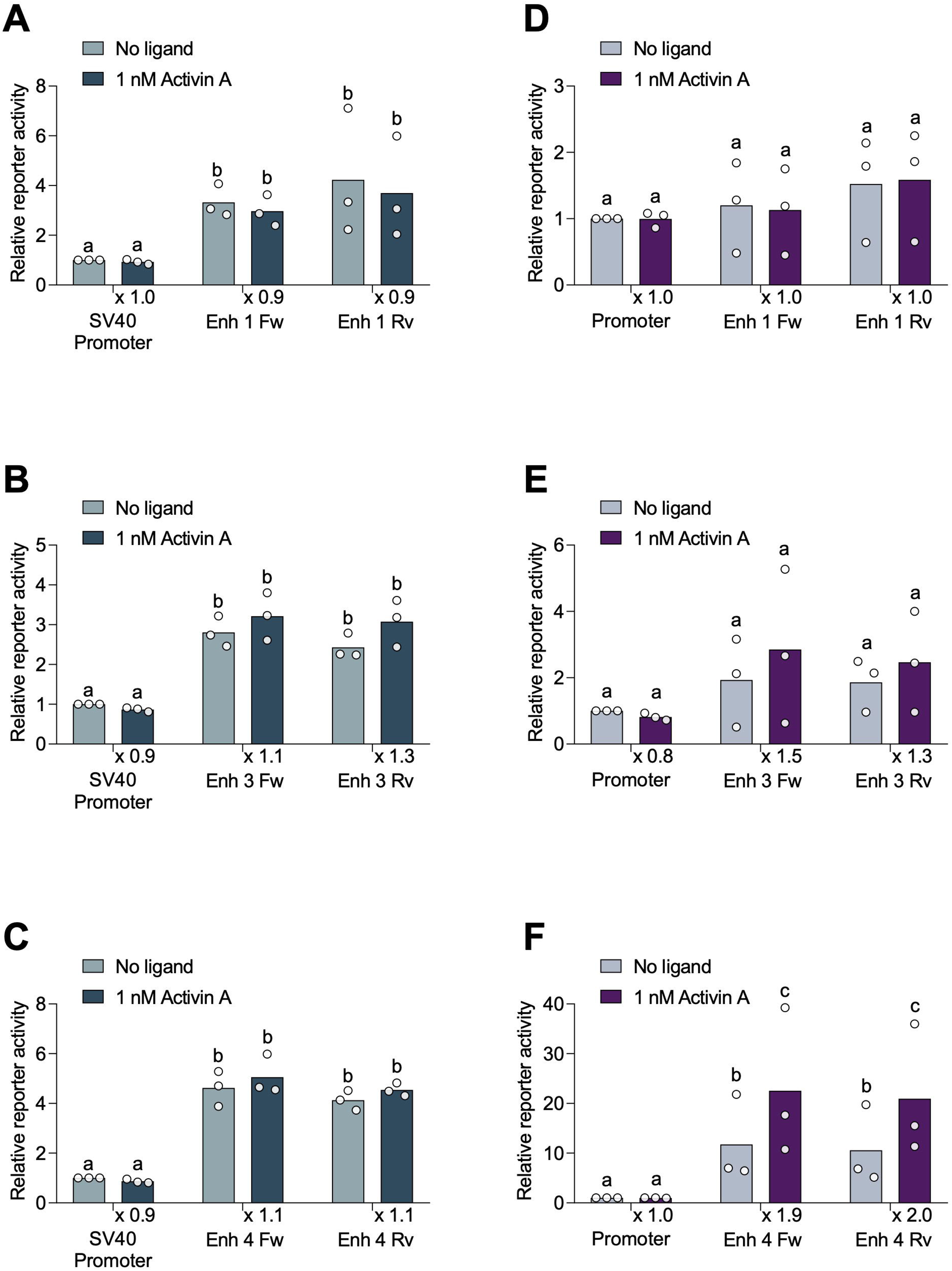
Enh 1, 3, or 4 effects in heterologous promoter or cell line contexts. LβT2 cells were transfected with a *SV40* promoter-reporter vector without or with (A) Enh 1, (B) Enh 3, or (C) Enh 4 inserted in either orientation downstream of luciferase. Cells were treated with no ligand (faint bars) or 1 nM activin A (darker bars) for 6h. Fold-induction by activin A is shown below the x-axis. αT3-1 cells were transfected with the -1990/+1 murine *Fshb* promoter-reporter vector without or with (D) Enh 1, (E) Enh 3, or (F) Enh 4 inserted in either orientation downstream of luciferase. Cells were treated with no ligand (faint purple bars) or 1 nM activin A (darker purple bars) for 6h. Fold-induction by activin A is shown below x-axis. Individual points are independent experiments. Data were analyzed as described in Fig. 4. Bars with different letters differ significantly.

### Enh 4 increases *Fshb* promoter-reporter activity in αT3-1 cells

In contrast to LβT2 cells, the less mature gonadotrope-like cell line, αT3-1, does not express *Fshb* mRNA basally or in response to activins (62). Moreover, murine *Fshb* promoter-reporters are not stimulated by activin A in these cells (**Figs. 5D-F**) (20). We nevertheless asked whether Enh 1, 3, or 4 might possess enhancer activity in these cells. Neither Enh 1 nor Enh 3 significantly altered basal or activin A-stimulated murine *Fshb* promoter-reporter activity in αT3-1 cells (**Figs. 5D-E**). In contrast, Enh 4 increased basal activity and, for the first time in our experience, conferred activin A sensitivity to the murine *Fshb* promoter in this cell line (**Fig. 5F**).

### Enh 3 and 4 exhibit enhancer characteristics in LβT2 cells

To further determine whether and how these open chromatin regions function as enhancers, we characterized their properties in LβT2 cells. *Fshb* expression and FSH secretion are low in these cells but can be induced by activin A (60,63). We previously reported that the *Fshb* promoter and gene are compacted in LβT2 cells (36). Here, we observed that the regions corresponding to Enh 3 and 4 in gonadotropes in vivo (**Fig. 6(a)**) were open in this cell line basally and may become more accessible following 6 h activin A treatment (**Fig. 6(b-c)**). The *Fshb* gene, promoter, Enh 1, and Enh 2 remained compacted basally or in the presence of activin A.

**Figure 6.**
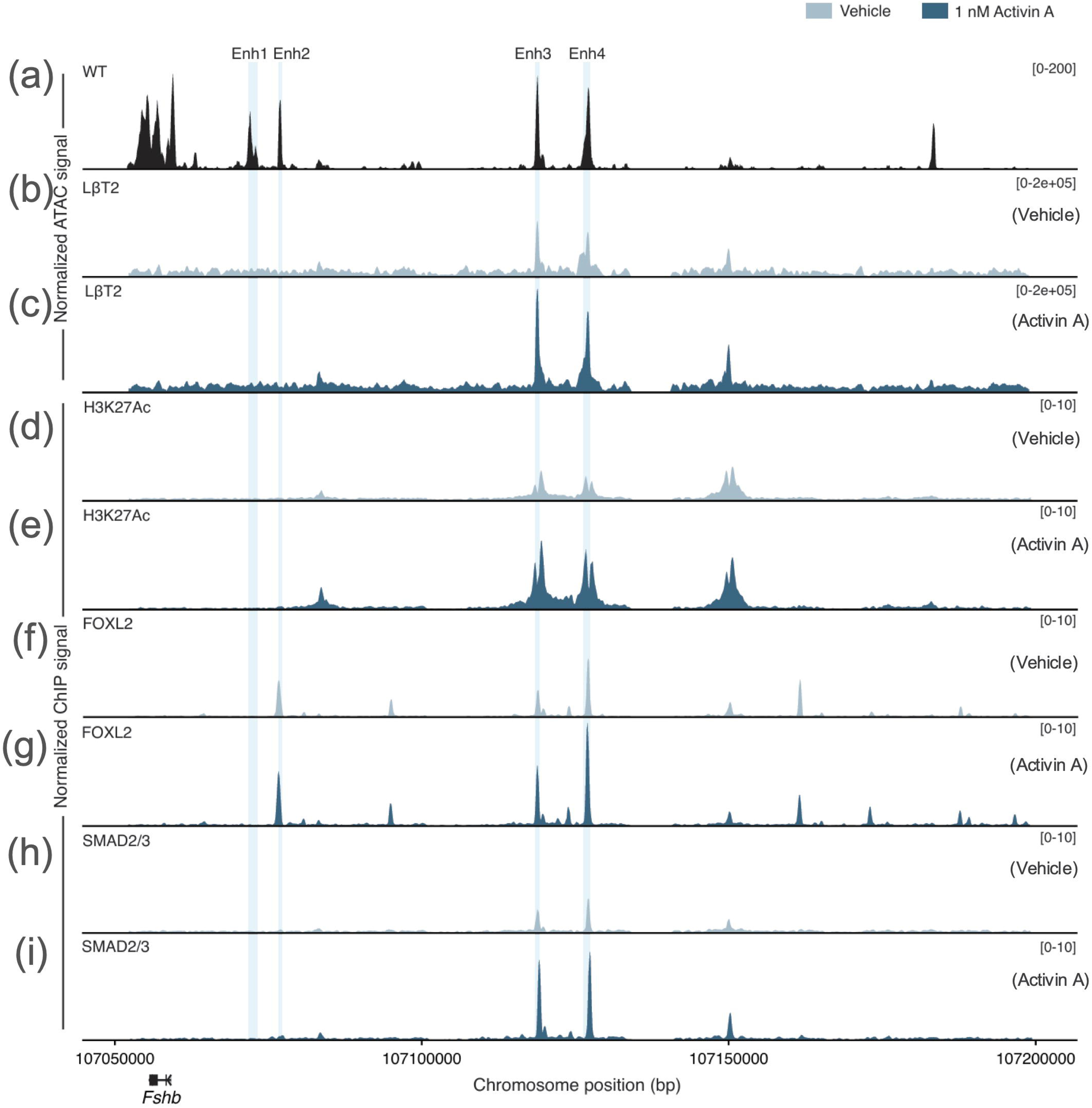
Enh 3 and 4 are open and enriched with H3K27Ac, FOXL2, and SMAD3 under basal and activin A stimulated conditions in LβT2 cells. (a) ATAC-seq data from WT mice in Fig. 3A(a) are aligned with bulk ATAC-seq in LβT2 cells treated with (b) vehicle or (c) 1 nM activin A for 6 h. ChIP-seq data showing enrichment of (d-e) H3K27Ac, (f-g) FOXL2, and (h-i) SMAD2/3 near the *Fshb* locus in LβT2 cells treated with (d, f, h) vehicle or (e, g, i) 1 nM activin A for 1 h.

In complementary ChIP-seq analyses, we observed enrichment of the enhancer mark, acetylated histone H3 at position K27 (H3K27Ac), at Enh 3 and Enh 4. This enrichment was increased by activin A (**Fig. 6(d-e)**). H3K27Ac was not observed at Enh 1 or Enh 2 basally or in response to activin A in these cells.

### SMADs and FOXL2 bind to *Fshb* enhancers 3 and 4 in LβT2 cells

Activin-stimulated *Fshb* transcription depends on binding of SMAD3, SMAD4, and FOXL2 to the promoter as revealed by reporter assays (12,21,23,25). Activin-dependent SMAD and FOXL2 binding to the *Fshb* promoter has been more difficult to demonstrate in native chromatin in LβT2 cells, perhaps because of its compacted state (**Fig. 6(b-c)**). We nevertheless asked whether activin A might promote SMAD and/or FOXL2 binding to the newly identified open chromatin regions. Antibodies for FOXL2 produce inconsistent results in ChIP in our experience (data not shown). Therefore, we generated LβT2 cells stably expressing a FLAG-tagged form of FOXL2 for these experiments. FLAG ChIP-seq (for FOXL2) revealed peaks of enrichment within Enh 2, Enh 3, and Enh 4, but not Enh 1 or the *Fshb* promoter (**Fig. 6(f-g)**). Activin A increased the FLAG-FOXL2 peak heights in Enh 2, Enh 3 and Enh 4. ChIP-seq similarly showed enrichment of SMAD2/3 in Enh 3 and Enh 4, which was enhanced by activin A (**Fig. 6(h-i)**). SMAD2/3 enrichment was not observed in the *Fshb* promoter, Enh 1, or Enh 2 basally or in response to activin A.

### FOXL2 binding to Enh 4 regulates *Fshb* promoter-reporter activity in LβT2 cells

Scrutiny of the ChIP-seq data showed that SMAD2/3 and FOXL2 were co-enriched in three regions within both Enh 3 and Enh 4. We used these data to help define minimal enhancers for follow-up reporter analyses. We first generated a series of enhancer truncations that removed one or more of the candidate SMAD/FOXL2 binding elements (SBE/FBE) in Enh 4 (**Supplemental Fig. S3A** in (34)). A 189 bp fragment containing only the most distal of the three elements, which spanned chr2 107,127,002-107,127,191 (hereafter referred to as Enh 4 mini), fully recapitulated the effects of the full-length Enh 4 on basal and activin A-stimulated murine *Fshb* promoter-reporter activity in LβT2 cells (**Fig. 7A, Supplemental Fig. S3A** in (34)).

**Figure 7.**
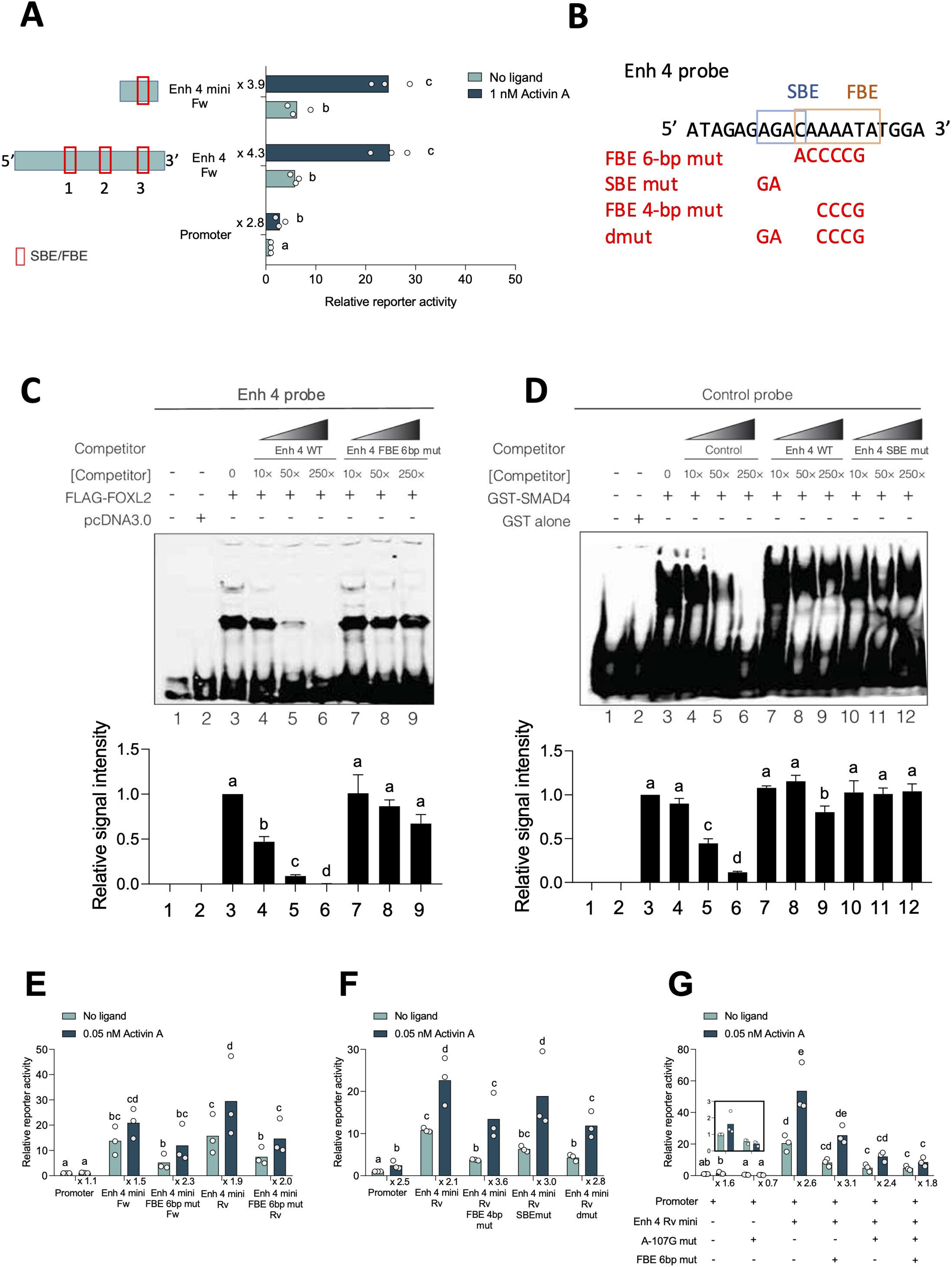
Binding of FOXL2 to Enh 4 confers enhancer activity. (A) LβT2 cells were transfected with -1990/+1 *Fshb* promoter-reporter vector alone, or containing the full-length, or minimal (mini) Enh 4 in the forward orientation downstream of luciferase. Positions of putative SBE/FBE sites are boxed in red. Cells were treated with no ligand (faint bars) or 1 nM activin (dark bars) for 6h. Fold induction by activin A is shown on the left of y-axis. (B) Sequence of the sense strand of a 20-bp Enh 4 probe. Positions of the putative SMAD and FOXL2 binding elements are boxed. Mutations used in competitor probes and reporters are labeled in red. (C) EMSA showing biotinylated Enh 4 probe incubated with nuclear extracts from HEK293T cells transfected with empty vector (pcDNA3.0, lane 2) or FLAG-FOXL2 (lane 3-9). Concentrations of the competitor probes in fold molar excess are labeled. The image is representative of four replicates of the experiment, which were quantified below. (D) EMSA using a biotinylated *Fshb* promoter probe (sequence shown in **Supplementary Table S1**) incubated with GST alone (lane 2) or recombinant GST-SMAD4-MH1 (lane 3-12). Competitor probes in fold molar excess were used as labeled. The image is representative of three independent experiments, which were quantified at the bottom of the panel. (E) LβT2 cells were transfected with the indicated reporters. The FBE mut refers to the 6-bp mutant. Cells were treated with vehicle (no ligand) or 0.05 nM activin A for 6h. Fold induction by activin A is shown below the x-axis for each reporter. (F) LβT2 cells were transfected with the indicated reporters. The FBEmut reflects the 4 bp mutant. Cells were treated with vehicle (no ligand) or 0.05 nM activin A for 6h. Fold induction by activin A is shown below the x-axis. (G) LβT2 cells were transfected with the indicated reporters. The FBE mut refers to the 6-bp mutant. Cells were treated with vehicle (no ligand) or 0.05 nM activin A for 6h. Fold induction by activin A is shown below the x-axis. Individual points are independent experiments. Bars with different letters differ significantly. Data were analyzed as described in Fig. 4. The inset shows the first two sets of bars at scale to facilitate their comparison.

Using electrophoretic mobility shift assays (EMSAs) with nuclear extracts from heterologous HEK293T cells over-expressing FLAG-tagged FOXL2, we identified base-pairs required for FOXL2 binding to a biotinylated double-stranded DNA probe (spanning chr2 107,127,140-107,127,160) containing the putative FOXL2 *cis*-element (FBE) in Enh 4 mini (**Figs. 7B-C**, lane 3). The binding was competed with an unlabeled wild-type probe but not with an unlabeled probe harboring 6 base-pair changes (Enh 4 FBE 6-bp mut) in the putative FBE (**Fig. 7C**, compare lanes 4-6 to 7-9; quantified at the bottom of the panel). In addition, an unlabeled probe harboring a 4-bp mutation in the FBE, which lacked a 1-bp overlap with an adjacent SBE (FBE 4-bp mut), similarly failed to compete for binding (**Fig. 7B** and data not shown).

SMAD3 can regulate *Fshb* transcription cooperatively rather than through direct DNA binding, as its MH1 domain (DNA binding domain) is not essential for *Fshb* expression in mice (21). Instead, SMAD4 primarily facilitates SMAD3/4 complex binding to the *Fshb* promoter (12,14). Therefore, we examined SMAD4 binding to Enh 4 using recombinant GST-SMAD4-MH1 in gel shifts. We observed only weak and inconsistent binding to the same probe used in the FOXL2 binding assays (data not shown). In contrast, GST-SMAD4-MH1 bound strongly to a positive control probe corresponding to the murine *Fshb* promoter (**Fig. 7D**, lane 3), as previously described (12). This binding could be competed efficiently by an unlabeled homologous promoter probe (lanes 4-6), but far less well by an unlabeled probe corresponding to Enh 4 (lanes 7-9; quantified at the bottom of the panel). Mutating the putative SBE in this probe eliminated the modest competitor activity (lanes 10-12). Similarly, the binding of GST-SMAD3-MH1 to the promoter probe could not be competed by the Enh 4 probe (**Supplemental Fig. S3B**, compare lanes 4-6 to 7-9 in (34)). Thus, direct SMAD4-MH1 or SMAD3-MH1 binding to Enh 4, if it occurs, appears to be weak.

We next interrogated the functional role(s) of the putative FBE and SBE in Enh 4 by introducing the same mutations used in the gel shift assays into reporter constructs. In initial experiments, the FBE mutation decreased basal, but not activin A-stimulated reporter activity (**Supplemental Fig. S3C** in (34)). Similar to full-length Enh 4 (**Fig. 4F**), the increase in basal reporter activity conferred by the minimal Enh 4 reflected endogenous activin-like signaling (**Supplemental Fig. S3D** in (34)) We speculated that the standard 1 nM activin A used in these assays might be saturating, thereby obscuring potential effects of the FBE mutation in Enh 4 on the response to exogenous activin A. Therefore, we performed a concentration-response experiment with the wild-type Enh 4 mini reporter, which revealed an EC50 of 0.05 nM for activin A (**Supplemental Fig. S3E** in (34)). The FBE mutation in Enh 4 mini reduced both basal and 0.05 nM activin A-stimulated reporter activity, though the fold activin A response was unaltered (**Fig. 7E**). The 4-bp FBE mutant, which lacks alterations to the putative SBE, behaved equivalently to the 6-bp FBE mutant in these assays (**Supplemental Fig. S3F** in (34)).

This latter result, combined with the weak binding activity in EMSAs, suggested that direct SMAD4 (or SMAD3) binding to Enh 4 might be dispensable. Indeed, when we introduced a 2-bp mutation in the SBE (**Fig. 7B** and **D**), there was a modest reduction in basal reporter activity, but no effect on the activin A response (**Fig. 7F**, fourth set of bars compared to second set of bars). The SBE mutation also had no further effect than that caused by the FBE mutation alone, as the reporter construct with both the SBE and FBE mutated (**Fig. 7B**, dmut) showed similar activity as the 4-bp FBE mut (**Fig. 7F**, fifth set of bars compared to third set of bars).

Thus, direct FOXL2 binding to Enh 4 appears to be more critical than direct SMAD binding for enhancer activity.

### Enh 4 activity also depends on the FOXL2 binding site in the *Fshb* promoter in LβT2 cells

Disrupting FOXL2 binding to the *Fshb* promoter with a point mutation at position -107 relative to the transcription start site (A-107G) inhibits induction by activin A in promoter-reporter assays (12,29). We next examined whether the effects of this mutation could be overcome or compensated by Enh 4 and FOXL2 binding therein. Activin A (at 0.05 nM) modestly stimulated murine *Fshb* promoter activity and this effect was abolished by the A-107G mutation (**Fig. 7G**, compare first two sets of bars; shown at higher magnification in the inset), as previously reported (12,29). Addition of the minimal Enh 4 again significantly increased basal and activin A-stimulated activity of the wild-type promoter (third set of bars). These effects were greatly attenuated, but not completely blocked, by the A-107G mutation (**Fig. 7G**, fifth set of bars). Mutating the FBE in Enh 4 (6-bp FBE mut) also reduced basal and activin A-stimulated reporter activity, though to a lesser extent than the promoter mutation (**Fig. 7G**, fourth set of bars). Mutating both FOXL2 binding sites (in the promoter and enhancer) had the most disruptive effects (**Fig. 7G**, last set of bars). The residual basal activity was completely abolished by blocking endogenous activin-like signaling with SB431542 (**Supplementary Fig. S3G** in (34)).

### SMAD4 binding to Enh 3 regulates *Fshb* promoter-reporter activity in LβT2 cells

Finally, we examined SMAD and FOXL2 binding and activity in Enh 3. We performed serial truncations, removing one or more of the candidate SBE/FBE in Enh 3 reporters. A minimal Enh 3 construct of 182 bp containing both putative SBE/FBE sites 1 and 2, spanning chr2 107,128,926 – 107,129,108, had higher enhancer activity than full-length Enh 3 in LβT2 cells (**Fig. 8A, Supplemental Fig. S4A** in (34)).

**Figure 8.**
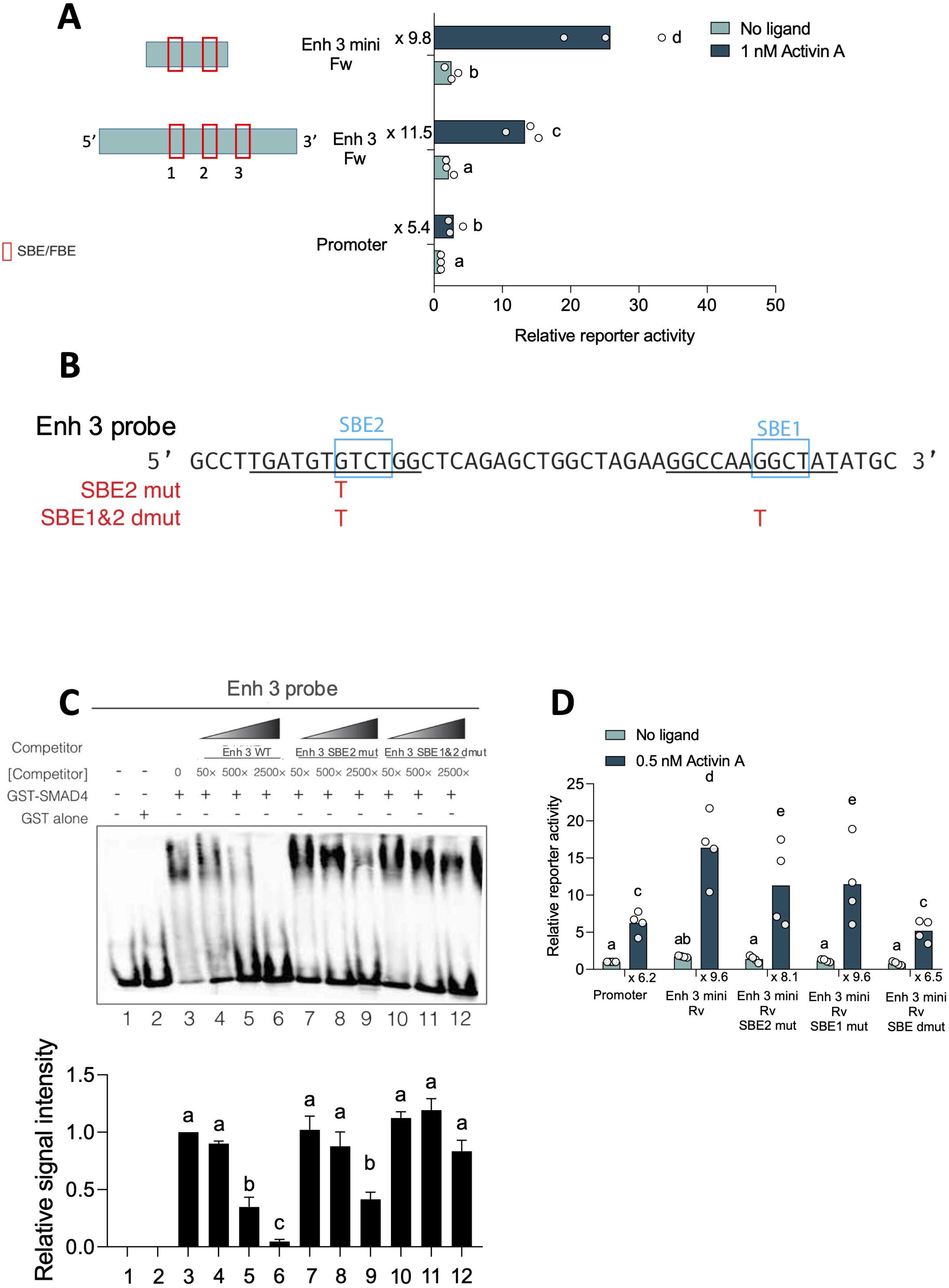
SMAD binding to Enh 3 confers enhancer activity. (A) LβT2 cells were transfected with the -1990/+1 *Fshb* promoter-reporter vector alone or containing the full-length or minimal (mini) Enh 3 in the forward orientation downstream of luciferase. The positions of putative SBE/FBE sites are boxed. Cells were treated with no ligand (faint bars) or 1 nM activin A (darker bars) for 6h. Fold induction by activin A is shown on the left side of the y-axis. (B) Sequence (sense strand only) of the 40-bp Enh 3 probe. Positions of SMAD binding elements are boxed. Mutations in competitor probes and reporters are marked in red. (C) Gel shift using biotinylated Enh 3 probe incubated with GST alone (lane 2) or recombinant GST-SMAD4-MH1 (lanes 3-12). Competitor probes at the indicated fold molar excess are labeled. The image is representative of four replicates of experiments, which were quantified below. (D) LβT2 cells were transfected with the indicated reporters. All enhancer constructs contained the minimal Enh 3 inserted in reverse orientation. Cells were treated with no ligand (faint bars) or 0.5 nM activin A (darker bars) for 6h. Fold induction by activin A is shown below the x-axis. Individual points are independent experiments. Data were analyzed as described in Fig. 4. Bars with different letters differ significantly.

In EMSAs, using nuclear extracts from heterologous cells over-expressing FOXL2 or purified GST-FOXL2-forkhead domain (FHD), we were unable to detect FOXL2 binding to a biotin-labeled Enh 3 probe spanning chr2 107,128,956 – 107,128,996, which contains both putative SBE/FBE 1 and 2 (data not shown). When used as a competitor, this probe also did not compete for binding of FOXL2 to the biotin-labeled Enh 4 probe used in the experiments in **Fig. 7C** (data not shown), suggesting that FOXL2 may not bind Enh 3 directly, at least on its own.

In contrast, and unlike what we observed with Enh 4, GST-SMAD4-MH1 or GST-SMAD3-MH1 bound to the biotin-labeled Enh 3 probe (**Fig. 8B-C**, lane 3, **Supplemental Fig. S4B**, lane 3 in (34)). This binding was competed with wild-type unlabeled Enh 3 probe (**Fig. 8C**, lanes 4-6, **Supplemental Fig. S4B,** lanes 4-6 in (34)), but less so with a probe containing a mutation in the second SBE (SBE2) (**Fig. 8C**, lanes 7-9, **Supplemental Fig. S4B,** lanes 7-9 in (34), sequence shown in **Fig. 8B**). Competition was completely abolished when the first SBE (SBE1) was also mutated (**Fig. 8C**, lanes 10-12, **Supplemental Fig. S4B,** lanes 10-12 in (34)). In addition, a shorter probe containing only SBE2 (**Supplemental Fig. S5A**) competed somewhat for binding **(Supplemental Fig. S5B**, lanes 7-9), yet a shorter probe containing SBE1 alone did not (**Supplemental Fig. S5B**, lanes 10-12 in (34)).

We then examined the effects of the SBE mutations in Enh 3 in reporter assays, using 0.5 nM activin A, as the 1 nM concentration might be saturating. Mutations in SBE1 or SBE2 had only minor effects on basal or activin A-stimulated reporter activity (**Fig. 8D**). Mutating both SBEs together, however, completely abrogated Enh 3 activity (**Fig. 8D**, compare the first and last sets of bars).

## Discussion

We report the existence of four open chromatin regions 5’ of *Fshb* that may function as transcriptional enhancers in mice. Three of these regions, as well as the *Fshb* promoter and gene, were compacted in gonadotropes of mice with genetic or pharmacologic blockade of activin type II receptor dependent signaling. When inserted downstream of luciferase in a murine *Fshb* promoter-luciferase plasmid, Enh 1, 3, or 4 significantly increased basal and/or activin A stimulated reporter activity. The increased basal activity, at least for Enh 3 and 4, reflected enhanced sensitivity to the actions of an endogenous TGFβ ligand, perhaps activin B. Enh 2, which was previously reported to act as a modest *Fshb* enhancer in LβT2 cells (30), lacked this activity in our experiments. It should be noted, however, that the constructs used in the two studies (both the length of the putative regulatory regions and their placement in the plasmids) were not identical. In addition, the heterogeneity of LβT2 cells (35) may also lead to differences in *Fshb* promoter-reporter regulation by Enh 2. Nevertheless, deletion of this enhancer region did not alter FSH production in mice, casting doubt on its necessity for *Fshb* expression in vivo.

During the preparation of this paper, normal FSH levels were similarly observed in a second mouse strain harboring a deletion of Enh 2 (64).

Enhancer redundancy is common (65). Knocking out Enh 2 did not affect chromatin accessibility of Enh 3 and 4 or the *Fshb* gene. The accessibility of Enh 1 may have been slightly reduced. Therefore, Enh 1, 3, and/or 4 may compensate for the loss of Enh 2 in KOs. This could be examined in mice harboring deletions of these putative enhancers in combination. Notably, as recently reported, deletion of a region encompassing both Enh 1 and 2 also failed to alter FSH production in mice (64). This, too, does not rule out the possibility for compensation by Enh 3 and/or 4. Indeed, these two regions had the greatest enhancer activity in our reporter assays, and they were uniquely in an open chromatin state and showed enrichment for H3K27Ac in LβT2 cells. Thus, Enh 3 and 4 may be particularly critical for FSH synthesis, at least in mice.

Our preliminary analysis of open chromatin regions upstream of *FSHB* in human gonadotropes (66) suggests that Enh 2 and 3, but not Enh 4, may be conserved (data not shown). It was previously reported that the human equivalent of Enh 2 has enhancer activity in reporter assays in LβT2 cells (30,31). This region also harbors two SNPs (rs11031005, rs11031006) associated with lower FSH levels in women (32,67). The major allele (G) for SNP rs11031006 is conserved in murine Enh 2 whereas the region containing the second SNP, rs11031005, is not present in mice. Substitution of the minor allele (A) for SNP rs11031006 did not affect FSH production in mice (64). Collectively, these data suggest that this enhancer may serve a more important role in humans than in mice.

Active enhancers and promoters can interact through chromatin looping (68,69). We did not investigate whether such looping occurs between the identified enhancers and the *Fshb* promoter in LβT2 cells because of the compaction of the latter in these cells (36). We therefore attempted to use 3C-methylation sequencing (70) to investigate looping in murine gonadotropes in vivo (data not shown). Unfortunately, the relatively low number of gonadotropes, even from 30 mice, rendered the assay insufficiently powered to obtain reliable results. Nevertheless, other data are consistent with the notion that such looping may occur. In particular, the same transcription factors that bind the *Fshb* promoter to mediate activin-dependent transcription in vitro (i.e., SMAD3, SMAD4, and FOXL2 (12,29)) also bind (either directly or indirectly) in Enh 3 and 4, and this binding was necessary for enhancer activity in reporter assays. Moreover, these proteins can homo- and heteromerized (14,21,23,29,71–74), suggesting that they may simultaneously bind both to the enhancers and promoter and provide a bridge between the regulatory regions. Indeed, binding of the same transcription factors in enhancers and promoters is common and can form the basis for chromatin looping (75–77).

Though the accessibility of the enhancers appears to be activin regulated in vivo, the available data suggest that activin sensitivity of *Fshb* is principally conferred by the promoter. First, Enh 1, 3 and 4 increased basal activity, but did not confer activin A responsiveness to the SV40 promoter in LβT2 cells. Second, addition of Enh 1, 3, or 4 increased overall activity of the *Fshb* promoter in LβT2 cells, but not the fold activin A response. Third, mutating the forkhead binding element (FBE) in Enh 4 or SMAD binding elements (SBEs) in Enh 3 were not as disruptive to the activin A response as was the FBE mutation in the *Fshb* promoter. Fourth, Enh 4 only mildly compensated for the mutation of the promoter FBE. Thus, the identified enhancers quantitatively increase transcription, whereas the *Fshb* promoter contains the critical regulatory sequences required for activin responsiveness.

The data may also shed some light on the activation of the *Fshb* locus developmentally. LβT2 cells are considered a ‘mature’ gonadotrope cell line, in that they express both gonadotropin β subunits. This contrasts with αT3-1 cells, which only express the common gonadotropin α subunit (78). Nevertheless, *Fshb* is expressed at far lower levels than *Lhb* in LβT2 cells (36,78). As we reported previously (36) and show again here, the *Fshb* promoter and gene are tightly compacted in these cells. Only Enh 3 and Enh 4 are accessible in the baseline state. LβT2 cells were derived from a tumor in a transgenic mouse in which the rat *Lhb* promoter was used to drive expression of the SV40 large T antigen (78). *Lhb* is reportedly expressed 1 day earlier than *Fshb* during murine pituitary development (79). Therefore, it is possible that the gonadotropes in this transgenic mouse began to transform before the cells were fully differentiated or the *Fshb* locus was fully open. This would suggest that Enh 3 and 4 may open in advance of the *Fshb* promoter, gene, and Enh 1 and 2. The mechanisms driving this chromatin state in LβT2 cells are not yet clear but may involve FOXL2. In fact, FOXL2 has demonstrated pioneer-like activity, as it binds predominantly to unopened chromatin regions in mouse embryonic stem cells, leading to increased local chromatin accessibility (80). Consistent with this idea, we also observed that in addition to Enh 3 and 4, FOXL2 was also associated with Enh 2, despite its closed conformation. Thus, FOXL2 may be necessary, but not sufficient to open *Fshb* enhancers and the promoter. Indeed, αT3-1 cells express FOXL2 but not *Fshb* (78,81).

However, when placed in close proximity, Enh 4 also increased both basal and activin-stimulated *Fshb* promoter reporter activity in αT3-1 cells. This demonstrates that all necessary trans-acting factors for Enh 4 function are already present and functional but may not be able to access the native locus, likely due to inactive chromatin states.

While ChIP-seq demonstrated co-occupancy of both FOXL2 and SMADs at Enh 3 and 4 in LβT2 cells, the binding may not be direct. EMSAs confirmed direct DNA binding only for FOXL2 to Enh 4 and SMADs to Enh 3, while reporter assays established the functional role of FOXL2 binding to Enh 4 and SMADs to Enh 3. The apparent discrepancy between genomic co-localization (by ChIP-seq) and the direct binding and functional data suggests that SMADs (likely SMAD4) may associate with Enh 4 and FOXL2 with Enh 3 through tethering mechanisms rather than canonical sequence-specific DNA binding. Such tethering could occur via protein-protein interactions with other factors (e.g., FOXL2 recruiting SMADs to Enh 4, or vice versa to Enh 3).

In summary, we identified four enhancers for murine *Fshb*, three of which were not previously described or characterized. The accessibility of three of the enhancers, like the *Fshb* promoter and gene, is dependent on signaling by TGFβ ligands through the activin type II receptors. In vitro, three of the enhancers increased *Fshb* promoter-reporter activity in LβT2 cells. In these cells, the most distal enhancers (Enh 3 and 4) were accessible and bound SMADs and FOXL2. These proteins were previously shown to mediate activin regulation of *Fshb* promoters from mice and pigs (82,83). Enhancers 2 and 3 appear to be conserved in humans, and their activity, at least in reporter assays, should be explored. Though Enh 2 is dispensable for FSH production in mice, it may be more important in humans, as SNPs in this region are associated with differences in FSH levels and humans may have less enhancer redundancy than mice. Deletion of Enh 1 and 2 in mice does not appear to affect FSH production (64), suggesting compensatory roles for Enh 3 and 4, which should be explored, especially as these are the only enhancers open in LβT2 cells and Enh 3 remains accessible in vivo when activin type II receptor signaling is blocked. The role, if any, of these enhancers in GnRH stimulated *Fshb* transcription should also be examined, as mechanisms of GnRH regulated FSH synthesis remain unresolved.

## Acknowledgements

We acknowledge the New York Genome Center and the Cedars-Sinai Applied Genomics, Computation & Translational Core for sequencing services. This work was supported in part by Cedars-Sinai institutional support. This work was supported in part through the computational and data resources and staff expertise provided by Scientific Computing at the Icahn School of Medicine at Mount Sinai and at the Cedars-Sinai Applied Genomics, Computation & Translational Core and the Salk Institute Ravazi Newman Integrative, Genomics and Bioinformatics Core. We thank Alissa Blackler for her technical contribution to ChIP-seq experiments. We acknowledge the expert contributions of Sven Heinz, Director of the Salk Institute Next Gen Sequencing Core and Christopher Benner, Director of the Salk Bioinformatics Core (both currently at the Department of Medicine at the University of California, San Diego) for help with design and data analysis of ChIP-seq experiments. We thank the indicated individuals for providing the indicated reagents.

## Data Availability

Supplementary data can be found in the Figshare Repository: https://figshare.com/articles/figure/_/30311416(34).

All sequencing datasets generated in this study have been deposited in the Gene Expression Omnibus (GEO). ATAC-seq data from LβT2 cells are available under accession GSE310458. ChIP-seq data from LβT2 cells are available under accession GSE310459. Single-nucleus ATAC-seq data from mouse pituitaries are available under accession GSE310460. All other data supporting this study are included in the Supplementary Information or available from the corresponding author upon request.

## References

1. Kumar TR, Wang Y, Lu N, Matzuk MM. Follicle stimulating hormone is required for ovarian follicle maturation but not male fertility. Nat Genet. 1997;15(2):201–204.

2. Tapanainen JS, Aittomäki K, Min J, Vaskivuo T, Huhtaniemi IT. Men homozygous for an inactivating mutation of the follicle-stimulating hormone (FSH) receptor gene present variable suppression of spermatogenesis and fertility. Nat Genet. 1997;15(2):205–206.

3. Bernard DJ, Fortin J, Wang Y, Lamba P. Mechanisms of FSH synthesis: what we know, what we don’t, and why you should care. Fertility and Sterility. 2010;93(8):2465–2485.

4. Zeleznik AJ, Midgley AR, Jr., Reichert LE, Jr. Granulosa cell maturation in the rat: increased binding of human chorionic gonadotropin following treatment with follicle-stimulating hormone in vivo. Endocrinology. 1974;95(3):818–825.

5. Steinkampf MP, Mendelson CR, Simpson ER. Regulation by follicle-stimulating hormone of the synthesis of aromatase cytochrome P-450 in human granulosa cells. Mol Endocrinol. 1987;1(7):465–471.

6. Oduwole OO, Peltoketo H, Huhtaniemi IT. Role of Follicle-Stimulating Hormone in Spermatogenesis. Front Endocrinol (Lausanne). 2018;9:763.

7. Attardi B, Miklos J. Rapid stimulatory effect of activin-A on messenger RNA encoding the follicle-stimulating hormone beta-subunit in rat pituitary cell cultures. Mol Endocrinol. 1990;4(5):721–726.

8. Lofrano-Porto A, Barra GB, Giacomini LA, Nascimento PP, Latronico AC, Casulari LA, da Rocha Neves Fde A. Luteinizing hormone beta mutation and hypogonadism in men and women. N Engl J Med. 2007;357(9):897–904.

9. Ongaro L, Zhou X, Wang Y, Schultz H, Zhou Z, Buddle ERS, Brûlé E, Lin Y-F, Schang G, Hagg A, Castonguay R, Liu Y, Su GH, Seidah NG, Ray KC, Karp SJ, Boehm U, Ruf-Zamojski F, Sealfon SC, …, Bernard DJ. Muscle-derived myostatin is a major endocrine driver of follicle-stimulating hormone synthesis. Science. 2025;387(6731):329–336.

10. Alonso CAI, David CD, Toufaily C, Wang Y, Zhou X, Ongaro L, Nudelman G, Nair VD, Ruf-Zamojski F, Boehm U, Sealfon SC, Bernard DJ. Activating Transcription Factor 3 Stimulates Follicle-Stimulating Hormone-β Expression In Vitro But Is Dispensable for Follicle-Stimulating Hormone Production in Murine Gonadotropes In Vivo. Endocrinology. 2023;164(5).

11. Lamba P, Santos MM, Philips DP, Bernard DJ. Acute regulation of murine follicle-stimulating hormone β subunit transcription by activin A. Journal of Molecular Endocrinology. 2006;36(1):201–220.

12. Tran S, Lamba P, Wang Y, Bernard DJ. SMADs and FOXL2 synergistically regulate murine FSHbeta transcription via a conserved proximal promoter element. Mol Endocrinol. 2011;25(7):1170–1183.

13. Fortin J, Ongaro L, Li Y, Tran S, Lamba P, Wang Y, Zhou X, Bernard DJ. Minireview: Activin Signaling in Gonadotropes: What Does the FOX say… to the SMAD? Mol Endocrinol. 2015;29(7):963–977.

14. Li Y, Schang G, Wang Y, Zhou X, Levasseur A, Boyer A, Deng CX, Treier M, Boehm U, Boerboom D, Bernard DJ. Conditional Deletion of FOXL2 and SMAD4 in Gonadotropes of Adult Mice Causes Isolated FSH Deficiency. Endocrinology. 2018;159(7):2641–2655.

15. Fortin J, Boehm U, Deng CX, Treier M, Bernard DJ. Follicle-stimulating hormone synthesis and fertility depend on SMAD4 and FOXL2. FASEB J. 2014;28(8):3396–3410.

16. Schang G, Ongaro L, Schultz H, Wang Y, Zhou X, Brule E, Boehm U, Lee SJ, Bernard DJ. Murine FSH Production Depends on the Activin Type II Receptors ACVR2A and ACVR2B. Endocrinology. 2020;161(7).

17. Thompson TB, Woodruff TK, Jardetzky TS. Structures of an ActRIIB:activin A complex reveal a novel binding mode for TGF-beta ligand:receptor interactions. Embo j. 2003;22(7):1555–1566.

18. Mathews LS, Vale WW. Expression cloning of an activin receptor, a predicted transmembrane serine kinase. Cell. 1991;65(6):973–982.

19. Ongaro L, Zhou X, Wang Y, Zhou Z, Schultz H, Buddle ERS, Brûlé E, Lin Y-F, Schang G, Castonguay R, Liu Y, Su GH, Seidah N, Ray KC, Karp SJ, Boehm U, Lee S-J, Bernard DJ. Myostatin is a major endocrine driver of follicle-stimulating hormone synthesis. bioRxiv. 2023:2023.2008.2030.555595.

20. Bernard DJ. Both SMAD2 and SMAD3 mediate activin-stimulated expression of the follicle-stimulating hormone beta subunit in mouse gonadotrope cells. Mol Endocrinol. 2004;18(3):606–623.

21. Li Y, Schang G, Boehm U, Deng CX, Graff J, Bernard DJ. SMAD3 Regulates Follicle-stimulating Hormone Synthesis by Pituitary Gonadotrope Cells in Vivo. J Biol Chem. 2017;292(6):2301–2314.

22. Suszko MI, Balkin DM, Chen Y, Woodruff TK. Smad3 mediates activin-induced transcription of follicle-stimulating hormone beta-subunit gene. Mol Endocrinol. 2005;19(7):1849–1858.

23. Lamba P, Wang Y, Tran S, Ouspenskaia T, Libasci V, Hebert TE, Miller GJ, Bernard DJ. Activin A regulates porcine follicle-stimulating hormone beta-subunit transcription via cooperative actions of SMADs and FOXL2. Endocrinology. 2010;151(11):5456–5467.

24. Suszko MI, Lo DJ, Suh H, Camper SA, Woodruff TK. Regulation of the rat follicle-stimulating hormone beta-subunit promoter by activin. Mol Endocrinol. 2003;17(3):318–332.

25. Corpuz PS, Lindaman LL, Mellon PL, Coss D. FoxL2 Is required for activin induction of the mouse and human follicle-stimulating hormone beta-subunit genes. Mol Endocrinol. 2010;24(5):1037–1051.

26. Tran S, Zhou X, Lafleur C, Calderon MJ, Ellsworth BS, Kimmins S, Boehm U, Treier M, Boerboom D, Bernard DJ. Impaired Fertility and FSH Synthesis in Gonadotrope-Specific Foxl2 Knockout Mice. Molecular Endocrinology. 2013;27(3):407–421.

27. Ongaro L, Schang G, Zhou Z, Kumar TR, Treier M, Deng C-X, Boehm U, Bernard DJ. Human Follicle-Stimulating Hormone ß Subunit Expression Depends on FOXL2 and SMAD4. Endocrinology. 2020;161(5).

28. Ghochani Y, Saini JK, Mellon PL, Thackray VG. FOXL2 is involved in the synergy between activin and progestins on the follicle-stimulating hormone β-subunit promoter. Endocrinology. 2012;153(4):2023–2033.

29. Lamba P, Fortin J, Tran S, Wang Y, Bernard DJ. A novel role for the forkhead transcription factor FOXL2 in activin A-regulated follicle-stimulating hormone beta subunit transcription. Mol Endocrinol. 2009;23(7):1001–1013.

30. Bohaczuk SC, Thackray VG, Shen J, Skowronska-Krawczyk D, Mellon PL. FSHB Transcription is Regulated by a Novel 5’ Distal Enhancer With a Fertility-Associated Single Nucleotide Polymorphism. Endocrinology. 2021;162(1).

31. Bohaczuk SC, Cassin J, Slaiwa TI, Thackray VG, Mellon PL. Distal Enhancer Potentiates Activin- and GnRH-Induced Transcription of FSHB. Endocrinology. 2021;162(7).

32. Golovchenko I, Aizikovich B, Golovchenko O, Reshetnikov E, Churnosova M, Aristova I, Ponomarenko I, Churnosov M. Sex Hormone Candidate Gene Polymorphisms Are Associated with Endometriosis. Int J Mol Sci. 2022;23(22).

33. Mbarek H, Steinberg S, Nyholt Dale R, Gordon Scott D, Miller Michael B, McRae Allan F, Hottenga Jouke J, Day Felix R, Willemsen G, de Geus Eco J, Davies Gareth E, Martin Hilary C, Penninx Brenda W, Jansen R, McAloney K, Vink Jacqueline M, Kaprio J, Plomin R, Spector Tim D, …, Boomsma Dorret I. Identification of Common Genetic Variants Influencing Spontaneous Dizygotic Twinning and Female Fertility. The American Journal of Human Genetics. 2016;98(5):898–908.

34. Jin Y SH, Ongaro L, Isidro Alonso CA, Schang G, Zhou X, Zamojski M, Nudelman G, Mendelev N, Onuma S, Welt CK, Bilezikjian LM, Sealfon SC, Ruf-Zamojski F, Bernard DJ. Regulation of murine follicle-stimulating hormone β subunit transcription by newly identified enhancers. Figshare Digital Repository 2025.

35. Ruf-Zamojski F, Ge Y, Pincas H, Shan J, Song Y, Hines N, Kelley K, Montagna C, Nair P, Toufaily C, Bernard D, Mellon P, Nair V, Turgeon J, Sealfon S. Cytogenetic, Genomic, and Functional Characterization of Pituitary Gonadotrope Cell Lines. Journal of the Endocrine Society. 2019;3:902–920.

36. Ruf-Zamojski F, Fribourg M, Ge Y, Nair V, Pincas H, Zaslavsky E, Nudelman G, Tuminello SJ, Watanabe H, Turgeon JL, Sealfon SC. Regulatory Architecture of the LβT2 Gonadotrope Cell Underlying the Response to Gonadotropin-Releasing Hormone. Front Endocrinol (Lausanne). 2018;9:34.

37. Stern E, Ruf-Zamojski F, Zalepa-King L, Pincas H, Choi SG, Peskin CS, Hayot F, Turgeon JL, Sealfon SC. Modeling and high-throughput experimental data uncover the mechanisms underlying Fshb gene sensitivity to gonadotropin-releasing hormone pulse frequency. J Biol Chem. 2017;292(23):9815–9829.

38. Buenrostro JD, Wu B, Chang HY, Greenleaf WJ. ATAC-seq: A Method for Assaying Chromatin Accessibility Genome-Wide. Curr Protoc Mol Biol. 2015;109:21.29.21–21.29.29.

39. Ewels PA, Peltzer A, Fillinger S, Patel H, Alneberg J, Wilm A, Garcia MU, Di Tommaso P, Nahnsen S. The nf-core framework for community-curated bioinformatics pipelines. Nat Biotechnol. 2020;38(3):276–278.

40. Langmead B, Salzberg SL. Fast gapped-read alignment with Bowtie 2. Nat Methods. 2012;9(4):357–359.

41. Stuart T, Srivastava A, Madad S, Lareau CA, Satija R. Single-cell chromatin state analysis with Signac. Nat Methods. 2021;18(11):1333–1341.

42. Ruf-Zamojski F, Zhang Z, Zamojski M, Smith GR, Mendelev N, Liu H, Nudelman G, Moriwaki M, Pincas H, Castanon RG, Nair VD, Seenarine N, Amper MAS, Zhou X, Ongaro L, Toufaily C, Schang G, Nery JR, Bartlett A, …, Sealfon SC. Single nucleus multi-omics regulatory landscape of the murine pituitary. Nat Commun. 2021;12(1):2677.

43. Mendelev N, Zamojski M, Amper MAS, Cheng WS, Pincas H, Nair VD, Zaslavsky E, Sealfon SC, Ruf-Zamojski F. Multi-omics profiling of single nuclei from frozen archived postmortem human pituitary tissue. STAR Protoc. 2022;3(2):101446.

44. Hao Y, Stuart T, Kowalski MH, Choudhary S, Hoffman P, Hartman A, Srivastava A, Molla G, Madad S, Fernandez-Granda C, Satija R. Dictionary learning for integrative, multimodal and scalable single-cell analysis. Nat Biotechnol. 2024;42(2):293–304.

45. Korsunsky I, Millard N, Fan J, Slowikowski K, Zhang F, Wei K, Baglaenko Y, Brenner M, Loh PR, Raychaudhuri S. Fast, sensitive and accurate integration of single-cell data with Harmony. Nat Methods. 2019;16(12):1289–1296.

46. Ongaro L, Alonso CAI, Zhou X, Brule E, Li Y, Schang G, Parlow AF, Steyn F, Bernard DJ. Development of a Highly Sensitive ELISA for Measurement of FSH in Serum, Plasma, and Whole Blood in Mice. Endocrinology. 2021;162(4).

47. Bernard DJ, Ongaro L. The Ultrasensitive Luteinizing Hormone (LH) ELISA Gets a New Lease on Life. Endocrinology. 2022;163(10).

48. Zawel L, Dai JL, Buckhaults P, Zhou S, Kinzler KW, Vogelstein B, Kern SE. Human Smad3 and Smad4 are sequence-specific transcription activators. Mol Cell. 1998;1(4):611–617.

49. Ruf-Zamojski F, Ge Y, Pincas H, Shan J, Song Y, Hines N, Kelley K, Montagna C, Nair P, Toufaily C, Bernard DJ, Mellon PL, Nair V, Turgeon JL, Sealfon SC. Cytogenetic, Genomic, and Functional Characterization of Pituitary Gonadotrope Cell Lines. J Endocr Soc. 2019;3(5):902–920.

50. Wang Y, Libasci V, Bernard DJ. Activin A induction of FSHβ subunit transcription requires SMAD4 in immortalized gonadotropes. Journal of Molecular Endocrinology. 2010;44(6):349–362.

51. Balcioglu O, Heinz RE, Freeman DW, Gates BL, Hagos BM, Booker E, Mirzaei Mehrabad E, Diesen HT, Bhakta K, Ranganathan S, Kachi M, Leblanc M, Gray PC, Spike BT. CRIPTO antagonist ALK4(L75A)-Fc inhibits breast cancer cell plasticity and adaptation to stress. Breast Cancer Res. 2020;22(1):125.

52. Blount AL, Vaughan JM, Vale WW, Bilezikjian LM. A Smad-binding element in intron 1 participates in activin-dependent regulation of the follistatin gene. J Biol Chem. 2008;283(11):7016–7026.

53. Heinz S, Benner C, Spann N, Bertolino E, Lin YC, Laslo P, Cheng JX, Murre C, Singh H, Glass CK. Simple combinations of lineage-determining transcription factors prime cis-regulatory elements required for macrophage and B cell identities. Mol Cell. 2010;38(4):576–589.

54. Huynh TN, Ren X. Producing GST-Cbx7 Fusion Proteins from Escherichia coli. Bio-protocol. 2017;7(12):e2333.

55. Zhen CY, Tatavosian R, Huynh TN, Duc HN, Das R, Kokotovic M, Grimm JB, Lavis LD, Lee J, Mejia FJ, Li Y, Yao T, Ren X. Live-cell single-molecule tracking reveals co-recognition of H3K27me3 and DNA targets polycomb Cbx7-PRC1 to chromatin. eLife. 2016;5:e17667.

56. Kvon EZ, Waymack R, Gad M, Wunderlich Z. Enhancer redundancy in development and disease. Nature Reviews Genetics. 2021;22(5):324–336.

57. Osterwalder M, Barozzi I, Tissières V, Fukuda-Yuzawa Y, Mannion BJ, Afzal SY, Lee EA, Zhu Y, Plajzer-Frick I, Pickle CS, Kato M, Garvin TH, Pham QT, Harrington AN, Akiyama JA, Afzal V, Lopez-Rios J, Dickel DE, Visel A, Pennacchio LA. Enhancer redundancy provides phenotypic robustness in mammalian development. Nature. 2018;554(7691):239–243.

58. Lach-Trifilieff E, Minetti GC, Sheppard K, Ibebunjo C, Feige JN, Hartmann S, Brachat S, Rivet H, Koelbing C, Morvan F, Hatakeyama S, Glass DJ. An antibody blocking activin type II receptors induces strong skeletal muscle hypertrophy and protects from atrophy. Mol Cell Biol. 2014;34(4):606–618.

59. Rejon CA, Ho CC, Wang Y, Zhou X, Bernard DJ, Hébert TE. Cycloheximide inhibits follicle-stimulating hormone β subunit transcription by blocking de novo synthesis of the labile activin type II receptor in gonadotrope cells. Cell Signal. 2013;25(6):1403–1412.

60. Pernasetti F, Vasilyev VV, Rosenberg SB, Bailey JS, Huang HJ, Miller WL, Mellon PL. Cell-specific transcriptional regulation of follicle-stimulating hormone-beta by activin and gonadotropin-releasing hormone in the LbetaT2 pituitary gonadotrope cell model. Endocrinology. 2001;142(6):2284–2295.

61. Inman GJ, Nicolás FJ, Callahan JF, Harling JD, Gaster LM, Reith AD, Laping NJ, Hill CS. SB-431542 is a potent and specific inhibitor of transforming growth factor-beta superfamily type I activin receptor-like kinase (ALK) receptors ALK4, ALK5, and ALK7. Mol Pharmacol. 2002;62(1):65–74.

62. Alarid ET, Windle JJ, Whyte DB, Mellon PL. Immortalization of pituitary cells at discrete stages of development by directed oncogenesis in transgenic mice. Development. 1996;122(10):3319–3329.

63. Graham KE, Nusser KD, Low MJ. LbetaT2 gonadotroph cells secrete follicle stimulating hormone (FSH) in response to active A. J Endocrinol. 1999;162(3):R1–5.

64. Bohaczuk SC, Tonsfeldt KJ, Slaiwa TI, Dunn GA, Gillette DLM, Yeo SE, Shi C, Cassin J, Thackray VG, Mellon PL. A Point Mutation in an Otherwise Dispensable Upstream Fshb Enhancer Moderately Impairs Fertility in Female Mice. Endocrinology. 2025;166(6).

65. Fletcher A WZ, Enciso G Shadow enhancers mediate trade-offs between transcriptional noise and fidelity. . PLoS Comput Biol 2023;19(5): e1011071.

66. Zhang Z, Zamojski M, Smith GR, Willis TL, Yianni V, Mendelev N, Pincas H, Seenarine N, Amper MAS, Vasoya M, Cheng WS, Zaslavsky E, Nair VD, Turgeon JL, Bernard DJ, Troyanskaya OG, Andoniadou CL, Sealfon SC, Ruf-Zamojski F. Single nucleus transcriptome and chromatin accessibility of postmortem human pituitaries reveal diverse stem cell regulatory mechanisms. Cell Rep. 2022;38(10):110467.

67. Hamdi M, Stacy S, Dale RN, Scott DG, Michael BM, Allan FM, Jouke Jan H, Felix RD, Gonneke W, Eco Jde G, Gareth ED, Hilary CM, Brenda WP, Rick J, Kerrie M, Jacqueline MV, Jaakko K, Robert P, Tim DS, …, Dorret IB. Identification of Common Genetic Variants Influencing Spontaneous Dizygotic Twinning and Female Fertility. The American Journal of Human Genetics. 2016;98(5):898–908.

68. Panigrahi A, O’Malley BW. Mechanisms of enhancer action: the known and the unknown. Genome Biology. 2021;22(1):108.

69. Popay TM, Dixon JR. Coming full circle: On the origin and evolution of the looping model for enhancer–promoter communication. Journal of Biological Chemistry. 2022;298(8):102117.

70. Lee DS, Luo C, Zhou J, Chandran S, Rivkin A, Bartlett A, Nery JR, Fitzpatrick C, O’Connor C, Dixon JR, Ecker JR. Simultaneous profiling of 3D genome structure and DNA methylation in single human cells. Nat Methods. 2019;16(10):999–1006.

71. Cheng JC, Klausen C, Leung PC. Overexpression of wild-type but not C134W mutant FOXL2 enhances GnRH-induced cell apoptosis by increasing GnRH receptor expression in human granulosa cell tumors. PLoS One. 2013;8(1):e55099.

72. Benayoun BA, Anttonen M, L’Hôte D, Bailly-Bechet M, Andersson N, Heikinheimo M, Veitia RA. Adult ovarian granulosa cell tumor transcriptomics: prevalence of FOXL2 target genes misregulation gives insights into the pathogenic mechanism of the p.Cys134Trp somatic mutation. Oncogene. 2013;32(22):2739–2746.

73. Kawabata M, Inoue H, Hanyu A, Imamura T, Miyazono K. Smad proteins exist as monomers *in vivo* and undergo homo- and hetero-oligomerization upon activation by serine/threonine kinase receptors. The EMBO Journal. 1998;17(14):4056–4065.

74. Blount AL, Schmidt K, Justice NJ, Vale WW, Fischer WH, Bilezikjian LM. FoxL2 and Smad3 coordinately regulate follistatin gene transcription. J Biol Chem. 2009;284(12):7631–7645.

75. Ray-Jones H, Spivakov M. Transcriptional enhancers and their communication with gene promoters. Cell Mol Life Sci. 2021;78(19-20):6453–6485.

76. Spitz F, Furlong EEM. Transcription factors: from enhancer binding to developmental control. Nature Reviews Genetics. 2012;13(9):613–626.

77. Vernimmen D, Bickmore WA. The Hierarchy of Transcriptional Activation: From Enhancer to Promoter. Trends Genet. 2015;31(12):696–708.

78. Alarid ET, Windle JJ, Whyte DB, Mellon PL. Immortalization of pituitary cells at discrete stages of development by directed oncogenesis in transgenic mice. Development. 1996;122(10):3319–3329.

79. Japón MA, Rubinstein M, Low MJ. In situ hybridization analysis of anterior pituitary hormone gene expression during fetal mouse development. J Histochem Cytochem. 1994;42(8):1117–1125.

80. Arora S, Yang J, Akiyama T, James DQ, Morrissey A, Blanda TR, Badjatia N, Lai WKM, Ko MSH, Pugh BF, Mahony S. Joint sequence & chromatin neural networks characterize the differential abilities of Forkhead transcription factors to engage inaccessible chromatin. bioRxiv. 2023.

81. Ellsworth BS, Egashira N, Haller JL, Butts DL, Cocquet J, Clay CM, Osamura RY, Camper SA. FOXL2 in the pituitary: molecular, genetic, and developmental analysis. Mol Endocrinol. 2006;20(11):2796–2805.

82. Tran S, Lamba P, Wang Y, Bernard DJ. SMADs and FOXL2 Synergistically Regulate Murine FSHβ Transcription Via a Conserved Proximal Promoter Element. Molecular Endocrinology. 2011;25(7):1170–1183.

83. Lamba P, Wang Y, Tran S, Ouspenskaia T, Libasci V, Hébert TE, Miller GJ, Bernard DJ. Activin A Regulates Porcine Follicle-Stimulating Hormone β-Subunit Transcription via Cooperative Actions of SMADs and FOXL2. Endocrinology. 2010;151(11):5456–5467.

