## Supplementary information for "Regulation of murine follicle-stimulating hormone β subunit transcription by newly identified enhancers"

#### Supplemental Figure Legends

**Supplemental Figure S1.** Generation of enhancer knockout mice. (A) Schematic representation of mice with the enhancer knocked out (chr2 107076832-107077562 (GRCm38/mm10)), using CRISPR-Cas9. Sequences of sgRNAs and the repair template are shown in Table 1. (B) Genotyping PCR of wild-type (WT) or enhancer knockout (KO) mice. Sequences of primers are shown in Table 1.

**Supplemental Figure S2.** Chromatin accessibility in and around the *Fshb* gene in the defined cell types of pituitaries of adult wild-type (A) male and (B) female mice.

**Supplemental Figure S3.** (A) LβT2 cells were transfected with the indicated reporters. All enhancer constructs are with Enh 4 truncations inserted in the forward orientation. Maps of relative locations of the SBE/FBE are shown at the left. Cells were treated with no ligand (faint bars) or 1 nM activin (dark bars) for 6h. Data were pooled from three different experiments. Dots of the same color indicate data points from the same experiment. (B) EMSA failing to show GST-SMAD3-MH1 binding to Enh 4. Biotinylated -132/-92 *Fshb* promoter probe (control probe, sequence shown in Table 1) was incubated with GST (lane 2) or recombinant GST-SMAD3-MH1 (lane 3-9). Competitor probes at the indicated fold molar excess were added as indicated. The image is representative of four replicate experiments, with the relative signal intensity of the four replicates plotted below. (C) LβT2 cells were transfected with the indicated reporters. Cells were treated with no ligand (faint bars) or 1 nM activin A (dark bars) for 6h. (D) LβT2 cells were transfected with the indicated reporters. Cells were treated with DMSO as vehicle (no ligand, faint bars) or 10 uM SB431542 (dark bars) overnight. (E) LβT2 cells were transfected with the Enh 4 mini reporter and treated with activin A at the indicated concentrations for 6h. (F) LβT2 cells were transfected with -1990/+1 *Fshb* promoter-reporter alone or with Enh 4 mini WT, Enh 4 mini 6-bp FBE mutant, or Enh 4 mini 4-bp FBE mutant. Cells were treated with either no ligand (faint bars) or 0.05 nM activin A (dark bars) for 6h. (G) LβT2 cells were transfected with the indicated reporters. Cells were treated with DMSO as vehicle (no ligand, faint grey bars) or 10 uM SB431542 (grey bars) overnight. Data were analyzed as described in Fig. 4. Bars with different letters differ significantly.

**Supplemental Figure S4.** (A) LβT2 cells were transfected with the indicated reporters. All Enh 3 constructs are with the enhancer inserted in the forward (fw) orientation. Cells were treated with no ligand (faint bars) or 1 nM activin A (dark bars) for 6h. Fold induction by activin A is shown at the left of the y-axis. Data were analyzed as described in Fig. 3. Different letters indicate bars that differ significantly. (B) EMSA showing binding of GST-SMAD3-MH1 to Enh 3 can be abrogated by the SBE2 mutation. Biotinylated Enh 3 probe (Fig. 8B) was incubated with GST (lane 2) or recombinant GST-SMAD3-MH1 (lane 3-12). Competitor probes were used at the indicated fold molar excess. The image is representative of three replicates, with the relative signal intensity of the three replicates plotted below.

**Supplemental Figure S5.** (A) Sequence (sense strand only) of the 40-bp Enh 3 wild-type (wt) probe containing both SBE site 1 and 2 as shown in Fig. 8B. Sequences of wt and mutant competitor probes are also shown. (B) Biotinylated wild-type Enh 3 probe was incubated with GST alone (lane 2) or recombinant GST-SMAD4-MH1 (lanes 3-15). Competitor probes and their relative molar concentrations are indicated. The image is representative of three experimental replicates. Quantification of all three assays is shown at the bottom. Bars with different letters differ significantly.

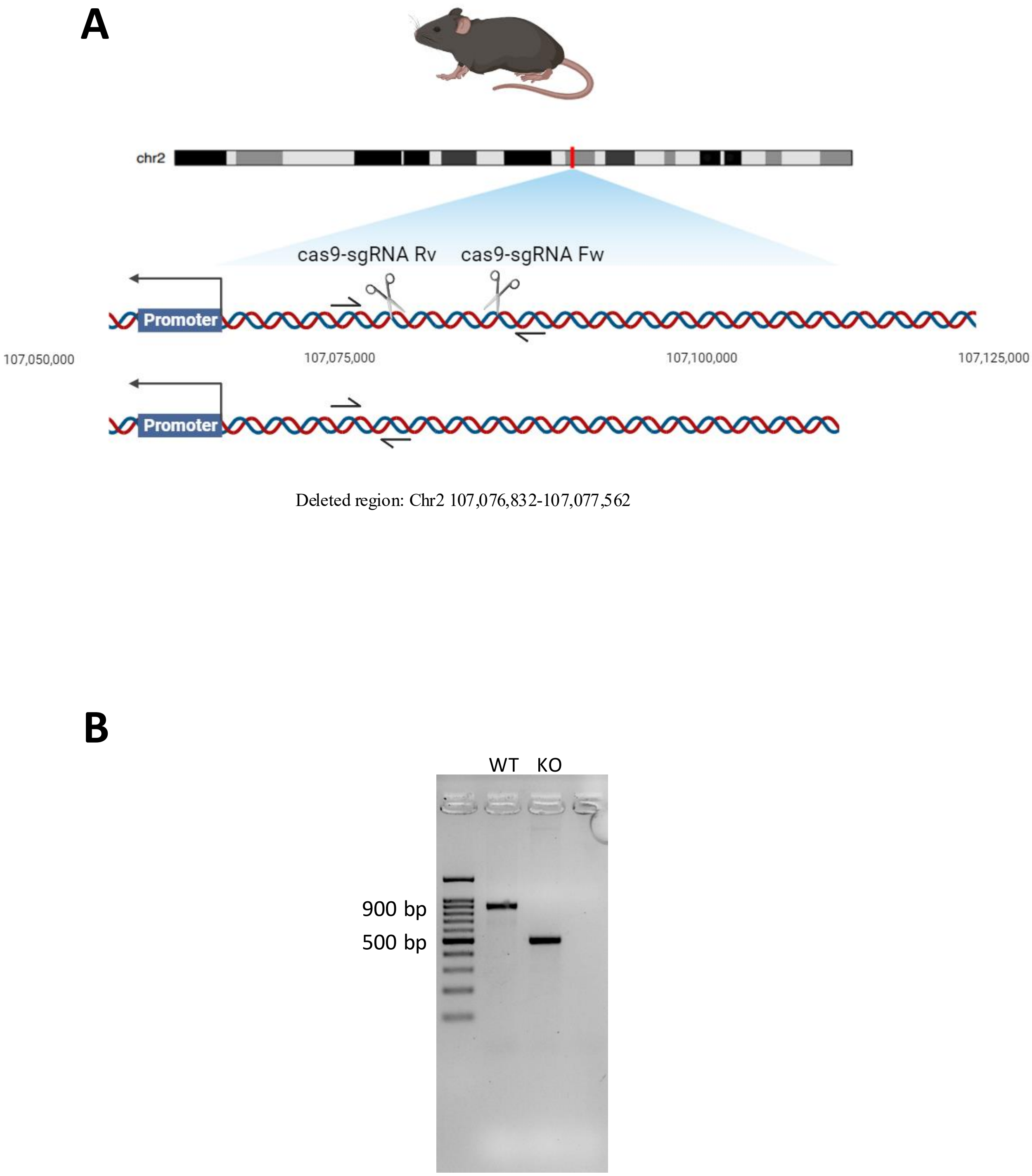
A
chr2
cas9-sgRNA Rv
cas9-sgRNA Fw
Promoter
107,050,000
107,075,000
107,100,000
107,125,000
Promoter
Deleted region: Chr2 107,076,832-107,077,562
B
WT
KO
900 bp
500 bp

Figure S1

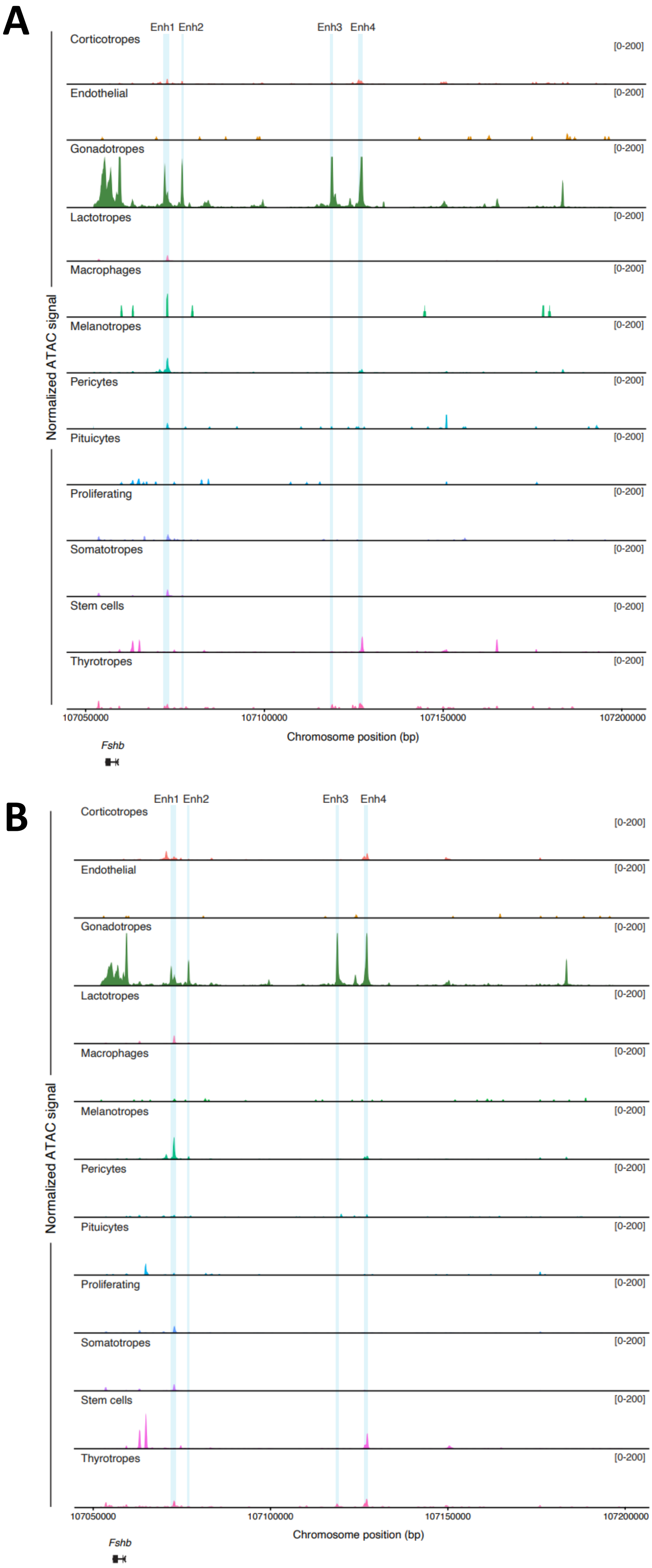
A
Enh1 Enh2
Enh3 Enh4
Normalized ATAC signal
Corticotropes
Endothelial
Gonadotropes
Lactotropes
Macrophages
Melanotropes
Pericytes
Pituicytes
Proliferating
Somatotropes
Stem cells
Thyrotropes
[0-200]
107050000
107100000
107150000
107200000
Chromosome position (bp)
*Fshb*
B
Enh1 Enh2
Enh3 Enh4
Normalized ATAC signal
Corticotropes
Endothelial
Gonadotropes
Lactotropes
Macrophages
Melanotropes
Pericytes
Pituicytes
Proliferating
Somatotropes
Stem cells
Thyrotropes
[0-200]
107050000
107100000
107150000
107200000
Chromosome position (bp)
*Fshb*

Figure S2

### Figure S3

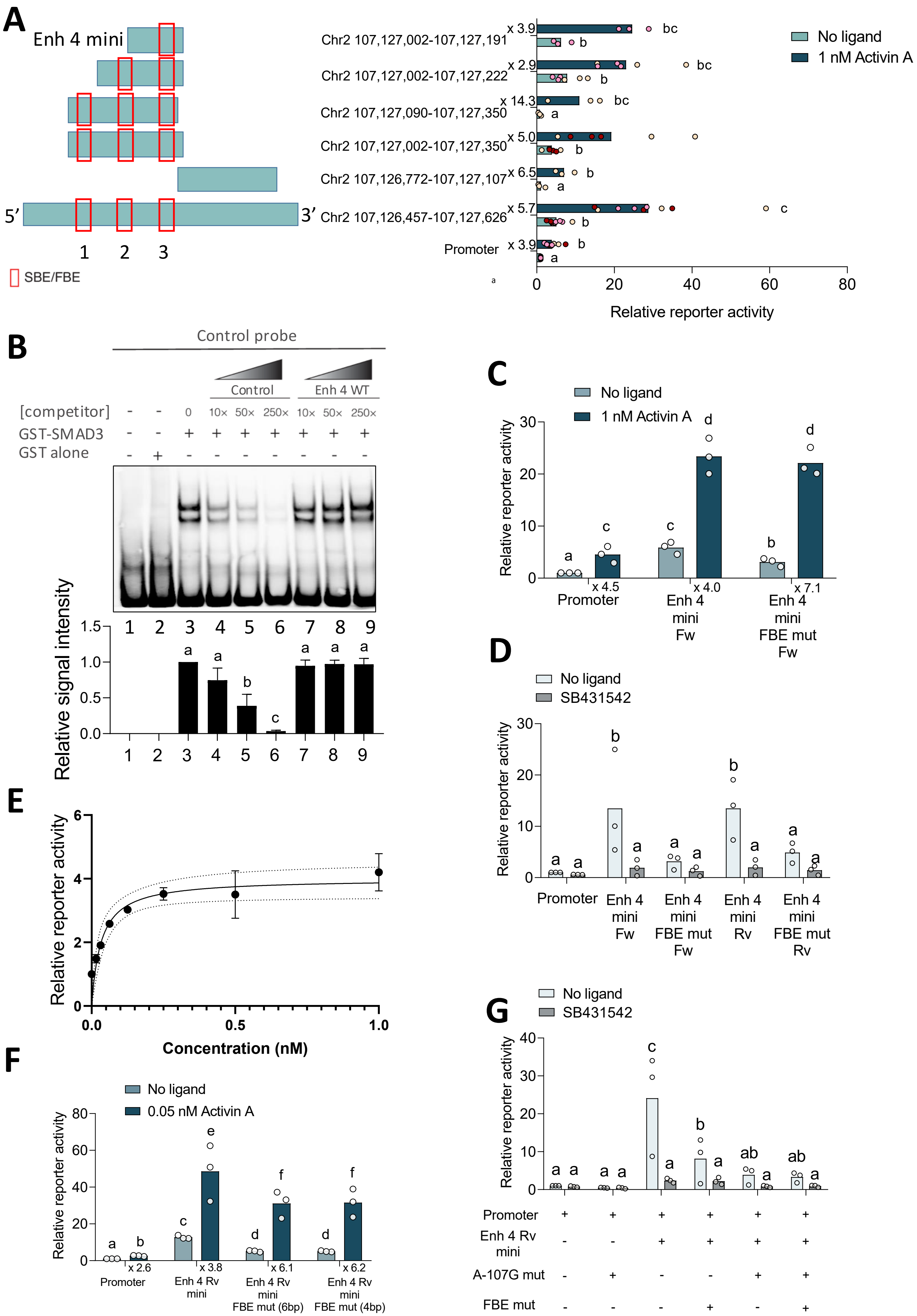

A
Enh 4 mini
5′
3′
1 2 3
SBE/FBE
Chr2 107,127,002-107,127,191
Chr2 107,127,002-107,127,222
Chr2 107,127,090-107,127,350
Chr2 107,127,002-107,127,350
Chr2 107,126,772-107,127,107
Chr2 107,126,457-107,127,626
Promoter
x 3.9
x 2.9
x 14.3
x 5.0
x 6.5
x 5.7
x 3.9
No ligand
1 nM Activin A
Relative reporter activity
B
Control probe
Control
Enh 4 WT
[competitor]
GST-SMAD3
GST alone
0 10× 50× 250× 10× 50× 250×
Relative signal intensity
C
No ligand
1 nM Activin A
Relative reporter activity
Promoter
Enh 4 mini Fw
Enh 4 mini FBE mut Fw
x 4.5
x 4.0
x 7.1
D
No ligand
SB431542
Relative reporter activity
Promoter
Enh 4 mini Fw
Enh 4 mini FBE mut Fw
Enh 4 mini Rv
Enh 4 mini FBE mut Rv
E
Relative reporter activity
Concentration (nM)
F
No ligand
0.05 nM Activin A
Relative reporter activity
Promoter
Enh 4 Rv mini
Enh 4 Rv mini FBE mut (6bp)
Enh 4 Rv mini FBE mut (4bp)
x 2.6
x 3.8
x 6.1
x 6.2
G
No ligand
SB431542
Relative reporter activity
Promoter
Enh 4 Rv mini
A-107G mut
FBE mut

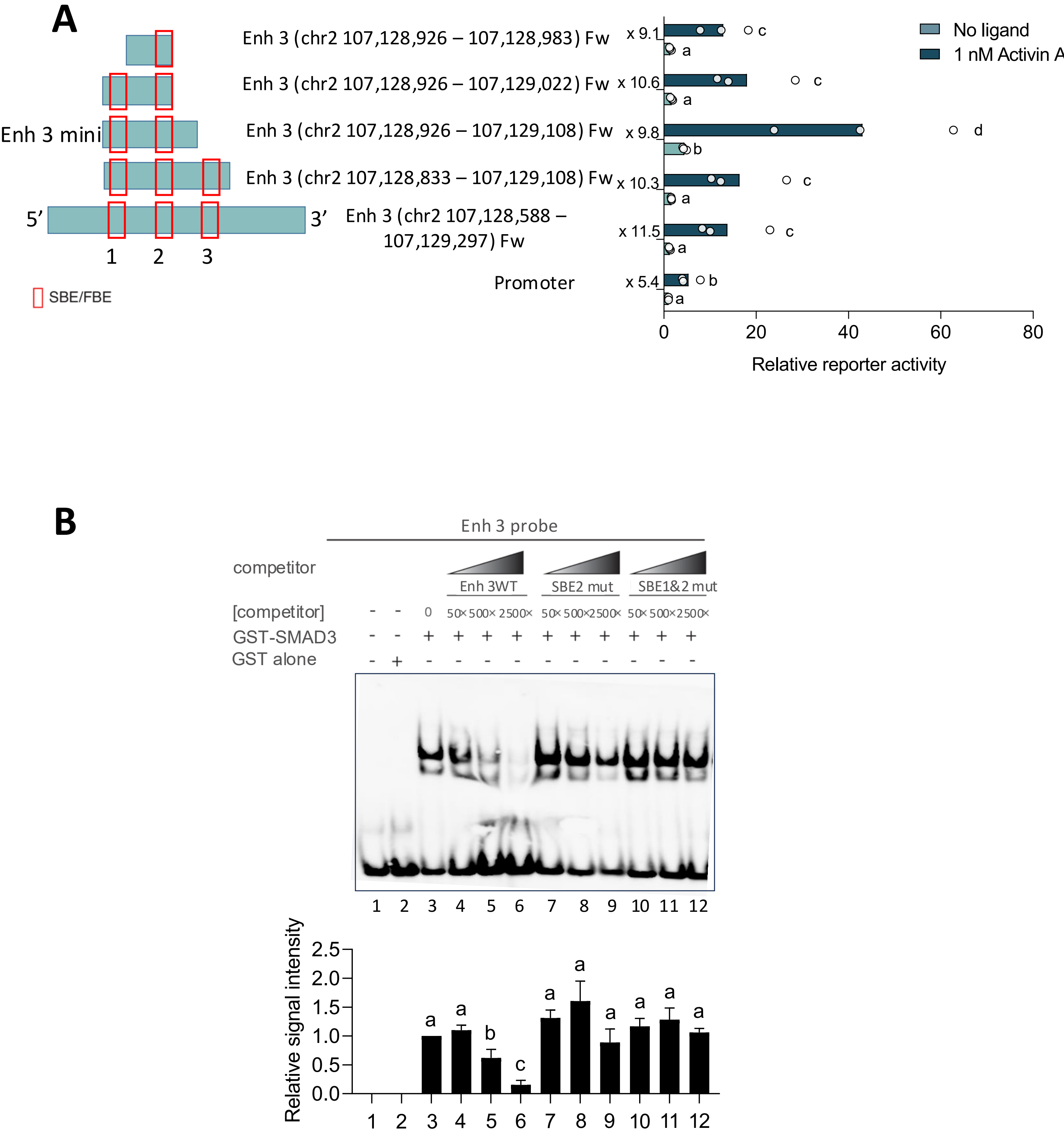
A
Enh 3 mini
5′
3′
1
2
3
SBE/FBE
Enh 3 (chr2 107,128,926 – 107,128,983) Fw
Enh 3 (chr2 107,128,926 – 107,129,022) Fw
Enh 3 (chr2 107,128,926 – 107,129,108) Fw
Enh 3 (chr2 107,128,833 – 107,129,108) Fw
Enh 3 (chr2 107,128,588 – 107,129,297) Fw
Promoter
x 9.1
x 10.6
x 9.8
x 10.3
x 11.5
x 5.4
No ligand
1 nM Activin A
Relative reporter activity
B
Enh 3 probe
competitor
Enh 3WT
SBE2 mut
SBE1&2 mut
[competitor]
GST-SMAD3
GST alone
Relative signal intensity

Figure S4

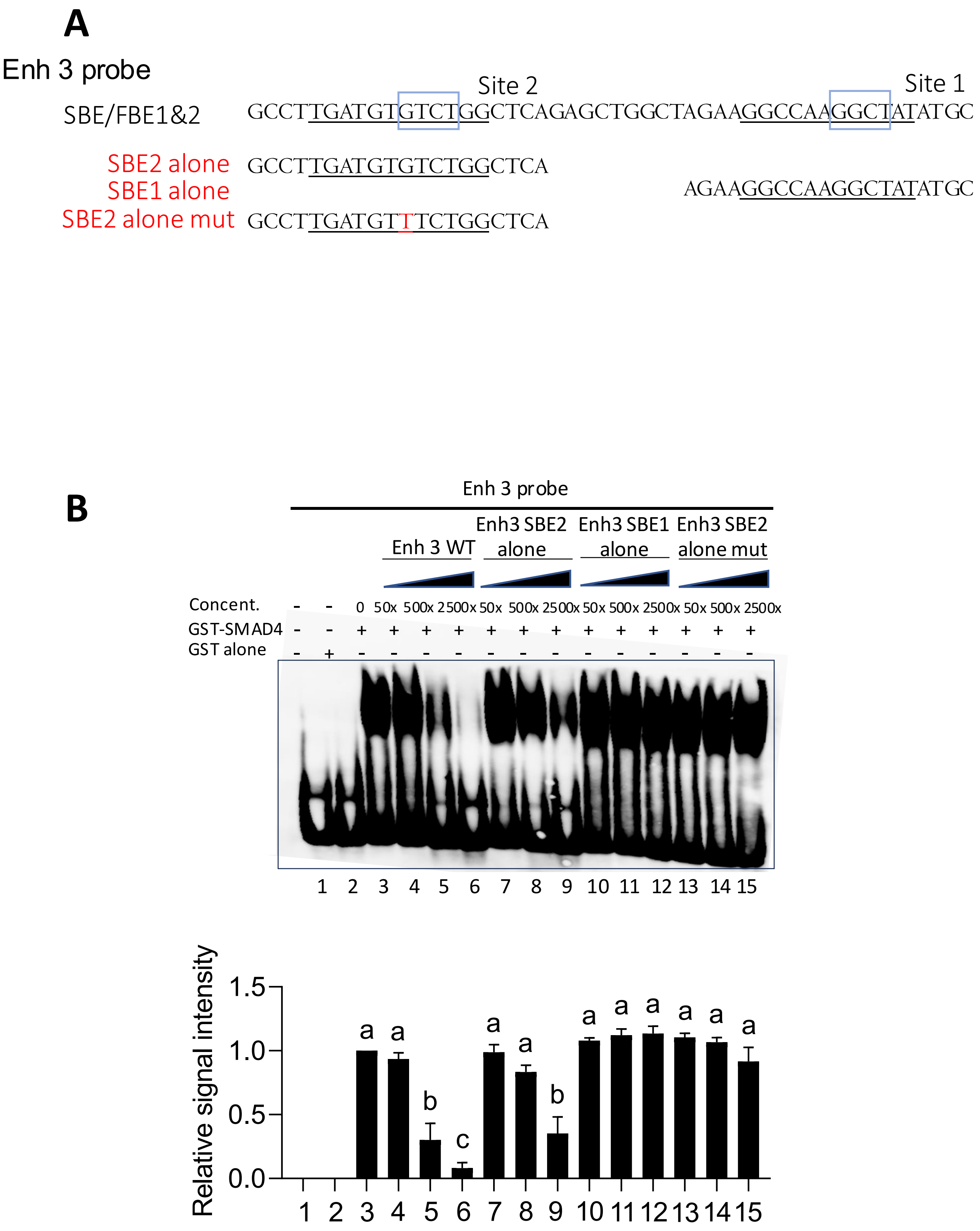
A
Enh 3 probe
Site 2
Site 1
SBE/FBE1&2 GCCTTGATGTGTCTGGCTCAGAGCTGGCTAGAAGGCCAAGGCTATATGC
SBE2 alone GCCTTGATGTGTCTGGCTCA
SBE1 alone AGAAGGCCAAGGCTATATGC
SBE2 alone mut GCCTTGATGTTTCTGGCTCA
B
Enh 3 probe
Enh 3 WT
Enh3 SBE2 alone
Enh3 SBE1 alone
Enh3 SBE2 alone mut
Concent. - - 0 50x 500x 2500x 50x 500x 2500x 50x 500x 2500x 50x 500x 2500x
GST-SMAD4 - - + + + + + + + + + + + + +
GST alone - + - - - - - - - - - - - - -
1 2 3 4 5 6 7 8 9 10 11 12 13 14 15
Relative signal intensity
1.5
1.0
0.5
0.0
a a b c a a b a a a a a a
1 2 3 4 5 6 7 8 9 10 11 12 13 14 15

Figure S5

**Supplemental Table S1. Primers and probes**

| | |
|---|---|
| **Single guide RNAs and repair template** | |
| sgRNA Fw | CCTGCTAGACATCAGGTACC |
| sgRNA Rv | GAGCATTAGATTCACTCTGC |
| repair template | TGTGTACCTGCATCTGCATTCTTCAGTAACGAAATTGACATCTGTTGCAAAGGGCACATGTACTCTTCAAGAAAATTTGATCTAGAATTTATCTTTGACATTCACTCTGCTGGTTTTGATGTTTAATAATACGCTGACTTTAACACTTGTGCATTTTTAATAATATAATATTAGACAATTATCACTTGATATTCTAATC |
| **Genotyping primers** | |
| Fw | GTGCCCATATGTCCCAAAGC |
| Rv | GGGTGTGGGACGTTACAGTG |
| **Cloning primers** | |
| Full length | |
| Enh 1 outer Fw | TCTGTTTTTAGGGCTGTGGCA |
| Enh 1 outer Rv | CATTGCCCCACATGGAATGC |
| Enh 1 inner_SalI Fw | AAAAGTCGACCCTGAGGCCAACCCTATGTT |
| Enh 1 inner_SalI Rv | AAAAGTCGACAGAAACCCAACTTTTACCTGTATGC |
| Enh 2 outer Fw | ACATGGTGATTTGAATGCCAAC |
| Enh 2 outer Rv | CCAAAGCTGAGGGGGATCTT |
| Enh 2 inner_SalI Fw | AAAAGTCGACTGCACAAGTGTTAAAGTCAGCG |
| Enh 2 inner_SalI Rv | AAAAGTCGACCATCTGTTGCAAAGGGCACA |
| Enh 3 outer Fw | AGGTTCCACATTCTGGGCTG |
| Enh 3 outer Rv | CCTTCAACCCCAACCACCTT |
| Enh 3 inner_SalI Fw | AAAAGTCGACAGCATCCTATTTAGAGCTGCTTG |
| Enh 3 inner_SalI Rv | AAAAGTCGACTGTGATGAGGAGAAGGCATCG |
| Enh 4 outer Fw | TGGCCTCCCATAGGTCCATT |
| Enh 4 outer Rv | AACGTCTTCATGATGGCCC |
| Enh 4 inner_SalI Fw | AAAAGTCGACTGGCCTCCCATAGGTCCATT |
| Enh 4 inner_SalI Rv | AAAAGTCGACGACTCCTTTGCCAAGATGCG |
| Truncations of Enh3 | |
| Enh 3 (245->) Fw | AAAAGTCGACCTGAGGCAGGCATTCTGTGT |
| Enh 3 (->520) Rv | AAAAGTCGACCAGTATACCAGGTCCTAGAGAAAGC |
| Enh 3 (338->) Fw | AAAAGTCGACACTGTGTATCCATAGCCTTGGG |
| Enh 3 (->434) Rv | AAAAGTCGACGCCAGCTCTGAGCCAGACA |
| Enh 3 (->395) Rv | AAAAGTCGACCTGTTTAGTCCTGGAGCAGCA |
| Truncations of Enh4 | |
| Enh 4 (315->) Fw | AAAAGTCGACTCTGTCTCCAGGTGGACCAA |
| Enh 4 (->650) Rv | AAAAGTCGACAATCGCACACAGCACAGA |

| | |
|---|---|
| Enh 4 (545->) Fw | AAAAGTCGACTCATTCTATTACATCTGTCTGGGAT |
| Enh 4 (->893) Rv | AAAAGTCGACTGTCTGAAAAGCAATCTGTAGCAA |
| Enh 4 (633->) Fw | AAAAGTCGACAATCGCACACAGCACAGACA |
| Enh 4 (->765) Rv | AAAAGTCGACGCTTTTCTGGAAATGAGCAGACT |
| Enh 4 (->734) Rv | AAAAGTCGACACACAAAGTGCCCTCCACTA |
| **Mutagenesis primers** | |
| Enh 4 FBE3 mut1 Fw | TCCACTAAGTCCCACTTTATCCATATTGGTTCTCTCTATAATAGCAACCTTGAATCC |
| Enh 4 FBE3 mut1 Rv | GGATTCAAGGTTGCTATTATAGAGAGAACCAATATGGATAAAGTGGGACTTAGTGGA |
| Enh 4 FBE3 mut2 Fw | CTCCACTAAGTCCCACTTTATCCATCGGGGTTCTCTCTATAATAGCAACCTTG |
| Enh 4 FBE3 mut2 Rv | CAAGGTTGCTATTATAGAGAGAACCCCGATGGATAAAGTGGGACTTAGTGGAG |
| mFshbp -A107G Fw | CTCCCTGTCCGTCTAAACGATGATTCCCTTTCAGCA |
| mFshbp -A107G Rv | TGCTGAAAGGGAATCATGTTTAGACGGACAGGGAG |
| Enh 4 SBE3 mut Fw | GGGATTCAAGGTTGCTATTATAGAGCTACAAAATATGGATAAAGTGGGACTT |
| Enh 4 SBE3 mut Rv | AAGTCCCACTTTATCCATATTTTGTAGCTCTATAATAGCAACCTTGAATCCC |
| Enh 4 FBESBE3 dmut Fw | AAGTCCCACTTTATCCATCGGGTGTAGCTCTATAATAGCAACCTTGAATCCC |
| Enh 4 FBESBE3 dmut Rv | GGGATTCAAGGTTGCTATTATAGAGCTACACCCGATGGATAAAGTGGGACTT |
| Enh 4 SBE revert back Fw | ACTTTATCCATCGGGTGTCTCTCTATAATAGCAACCTTGAATCCCT |
| Enh 4 SBE revert back Rv | AGGGATTCAAGGTTGCTATTATAGAGAGACACCCGATGGATAAAGT |
| Enh 3 SBE2 mut Fw | CTCTGAGCCAGAAACATCAAGGCCTTTTAGACCCAA |
| Enh 3 SBE2 mut Rv | TTGGGTCTAAAAGGCCTTGATGTTTCTGGCTCAGAG |
| Enh 3 SBE1 mut Fw | AGCAGCATATAGCATTGGCCTTCTAGCCAGCTC |
| Enh 3 SBE1 mut Rv | GAGCTGGCTAGAAGGCCAATGCTATATGCTGCT |
| **qPCR Primers** | |
| *Fshb* Fw | GTGCGGGCTACTGCTACACT |
| *Fshb* Rv | CAGGCAATCTTACGGTCTCG |
| *Lhb* Fw | ACTGTGCCGGCCTGTCAACG |
| *Lhb* Rv | AGCAGCCGGCAGTACTCGGA |
| *Cga* Fw | TCCCTCAAAAAGTCCAQGAGC |
| *Cga* Rv | GAAGAGAATGAAGAATATGCAG |
| *Gnrhr* Fw | TTCGCTACCTCCTTTGTCGT |
| *Gnrhr* Rv | CACGGGTTTAGGAAAGCAAA |

| | |
|---|---|
| *Rpl19* Fw | CGGGAATCCAAGAAGATTGA |
| *Rpl19* Rv | TTCAGCTTGTGGATGTGCTC |
| **EMSA probes** | |
| Enh 4 wt Sense | ATAGAGAGACAAAATATGGA |
| Enh 4 wt Antisense | TCCATATTTTGTCTCTCTAT |
| Enh 4 FBE mut Sense | ATAGAGAGAACCCCGATGGA |
| Enh 4 FBE mut Antisense | TCCATCGGGGTTCTCTCTAT |
| Enh 4 SBE mut Sense | ATAGAGCTACAAAATATGGA |
| Enh 4 SBE mut Antisense | TCCATATTTTGTAGCTCTAT |
| Enh 3 wt Sense | GCCTTGATGTGTCTGGCTCAGAGCTGGCTAGAAGGCCAAGGC TATATGC |
| Enh 3 wt Antisense | GCATATAGCCTTGGCCTTCTAGCCAGCTCTGAGCCAGACACAT CAAGGC |
| Enh 3 SBE2 mut Sense | GCCTTGATGTTTCTGGCTCAGAGCTGGCTAGAAGGCCAAGGC TATATGC |
| Enh 3 SBE2 mut Antisense | GCATATAGCCTTGGCCTTCTAGCCAGCTCTGAGCCAGAAACAT CAAGGC |
| Enh 3 SBE1&2 mut Sense | GCCTTGATGTTTCTGGCTCAGAGCTGGCTAGAAGGCCAATGCT ATATGC |
| Enh 3 SBE1&2 mut Antisense | GCATATAGCATTGGCCTTCTAGCCAGCTCTGAGCCAGAAACAT CAAGGC |
| Enh 3 SBE2 alone Sense | GCCTTGATGTGTCTGGCTCA |
| Enh 3 SBE2 alone Antisense | TGAGCCAGACACATCAAGGC |
| Enh 3 SBE1 alone Sense | AGAAGGCCAAGGCTATATGC |
| Enh 3 SBE1 alone Antisense | GCATATAGCCTTGGCCTTCT |
| Enh 3 SBE2 alone mut Sense | GCCTTGATGTTTCTGGCTCA |
| Enh 3 SBE2 alone mut Antisense | TGAGCCAGAAACATCAAGGC |

**Supplemental Table S2. Authentication of the LβT2 cell lines**

| | | LβT2b | LβT2 FOXL2 | LβT2 GFP | | |
|---|---|---|---|---|---|---|
| **Marker #** | **Chromosome** | **2** | **5** | **6** | **C57BL/6J** | **BALB/cJ** |
| MCA-4-2 | 4 | 20.3, 21.3 | 20.3, 21.3 | 20.3, 21.3 | 20.3 | 21.3 |
| MCA-5-5 | 5 | 13, 17 | 13, 17 | 13, 17 | 17 | 14 |
| MCA-6-4 | 6 | 19, 20 | 19, 20 | 19, 20 | 18 | 17 |
| MCA-6-7 | 6 | 12 | 12 | 12 | 17, 18 | 12 |
| MCA-9-2 | 9 | 18 | 18 | 18 | 18 | 15 |
| MCA-12-1 | 12 | 16, 17 | 16, 17 | 16, 17 | 17 | 16 |
| MCA-15-3 | 15 | 22.3 | 22.3 | 22.3 | 22.3 | 22.3 |
| MCA-18-3 | 18 | 16, 17 | 16, 17 | 16, 17 | 16 | 18 |
| MCA-X-1 | X | 25 | 25 | 25 | 27 | 24 |
| MCA-1-1 | 1 | 16,17 | 16,17 | 16,17 | 16 | 15 |
| MCA-1-2 | 1 | 19 | 19 | 19 | 18, 19 | 17 |
| MCA-2-1 | 2 | 16 | 16 | 16 | 16 | 16 |
| MCA-3-2 | 3 | 13 | 13 | 13 | 14 | 14 |
| MCA-7-1 | 7 | **26.2** | **26.2** | **26.2, 27.2** | 26.2 | 26.2, 27.2 |
| MCA-8-1 | 8 | 16 | 16 | 16 | 16 | 13 |
| MCA-11-2 | 11 | 16, 18 | 16, 18 | 16, 18 | 16 | 16, 17 |
| MCA-13-1 | 13 | 16.2 | 16.2 | 16.2 | 17 | 14.2 |
| MCA-17-2 | 17 | 15 | 15 | 15 | 15 | 16 |
| MCA-19-2 | 19 | 12 | 12 | 12 | 13 | 12, 13 |
